# Revision of Ambisporaceae, with three new genera and one new species and a morphological identification key for all the species currently attributed to this family

**DOI:** 10.64898/2026.02.11.705428

**Authors:** Gladstone Alves da Silva, Ewald Sieverding, Viviane Monique Santos, Claudia Castillo, Samar Velho da Silveira, Thays Gabrielle Lins de Oliveira, Daniele Magna Azevedo de Assis, Paulo Vitor Dutra de Souza, Mike Anderson Corazon-Guivin, Iván Sánchez-Castro, Javier Palenzuela, Fritz Oehl

## Abstract

The objective of this study was to re-analyse the molecular phylogeny and/or the morphology of all species, which have been attributed to the so-far mono-generic fungal family Ambisporaceae. The genus *Ambispora* has been well-known for its spore bi-morphy described even from single spore clusters. Triple-walled spores are differentiated on sporiferous saccules, while mono-walled spores are formed on simple subtending hyphae. New phylogenetic analyses reveal dissimilarities of ≥10% in partial nrDNA gene of three different stable phylogenetic clades and thus suggest the division of *Ambispora* into three genera, which simultaneously request for advanced morphological separations. These advances are primarily based on the more diverse spore wall composition of the ambisporoid-acaulosporoid morph rather than on the rather simple-glomoid morph. While all known species of the triple-walled morph have an evanescent to semi-permanent outer spore wall, i) *Am. fennica*, *Am. brasiliensis*, *Am. gerdemannii* and *Am. nicolsonii* have a smooth, permanent central spore wall (*Am. fennica* clade, A), ii) the central wall of *Am. appendicula*, *Am. callosa*, *Am. leptoticha* and *Am. jimgerdemannii* is alveolate (*Am. appendicula* clade, B), and iii) the central wall of *Am. granatensis* is smooth, but easily degraded, thus rather short-lived and not permanent but evanescent (*Am. granatensis* clade, C). In conclusion, species of the *Am. fennica* clade represent the genus *Ambispora*, while species of the *Am. appendicula* clade represent the new genus *Appendiculaspora*, and the mono-specific *Am. granatensis* clade represents the new genus *Ephemerapareta*. Species of an additional morph, with triple-walled spores, but apparently formed on subtending hyphae, and having a diagnostic reticulate, football-like middle wall, are here separated from the revised genus *Ambispora* based solely on morphological analyses, since molecular identification analyses so far failed and remained merely unknown. This later morph and genus is based on the type species *Pelotaspora reticulata* comb. nov, and on *P. austrolatina* sp. nov. Concomitant molecular phylogenetic and morphological analyses are needed to attribute not only *Pelotaspora* spp., but also those species, for which hitherto only the ambisporoid-glomoid morph has been observed correctly within the family Ambisporaceae. Without molecular analyses, such species with glomoid but unknown ambisporoid-acaulosporoid morph have to be retained within *Ambispora*.

## Introduction

The introduction of *Appendicispora*/*Ambispora* into the Archaeosporaceae was one of the most ‘lively’ coincidences in the history of AM fungal taxonomy (Spain et al., 2006; Walker et al., 2007a, 2007b, 2008). *Appendicispora* and *Ambispora* were described shortly one after the other, virtually for the same fungal clade, but based on two different type species (*Ap. appendicula* and *Am. fennica*, respectively (Spain et al., 2006; Walker et al., 2007a). Since, based on nomenclatural rules, *Appendicispora* (Spain et al., 2006) was a late homonym of *Appendicospora* (Ascomycota), finally *Ambispora* (Walker, 2008) was established as the legitim genus name, although it was described after *Appendicispora*.

Already the description of the first few species, at date attributed to the genus *Ambispora*, was literally confusing for several reasons. *Glomus gerdemannii*, currently *Ambispora gerdemannii*, was described based on its ambisporoid-acaulosporoid morph, which had erroneously been judged to be a typical glomoid morph of the genus *Glomus* (Rose et al., 1979). A second species, called *Acaulospora gerdemannii* (Nicolson and Schenck, 1979), was described by its ‘acaulosporoid’ morph, which did not fit with ambisporoid-acaulosporoid morph of *G. gerdemannii*, and neither with the acaulosporoid morph of *Ac appendicula* (= *Am. appendicula*; Schenck et al., 1984; Spain et al., 2006). Finally, besides *Am. appendicula*, for which both acaulosporoid and glomoid morphs had been described directly from single species isolates (Schenck et al., 1984), another species had been described, which was *G. leptotichum*, based on its glomoid morph and its diagnostic reticulate ornamentation on its outer spore surfaces (Schenck and Smith, 1982), which is not existing in the glomoid morph of *Am. appendicula* or *Am. gerdemannii*, and thus currently has been called *Am. leptoticha*. In between these years, *Ac. nicolsonii*, currently *Am. nicolsonii*, was described as a ‘typical’ *Acaulospora*, although the diagnostic beaded inner wall of Acaulosporaceae species has been missing, as in *Ac. gerdemannii* and *Ac. appendicula* (Nicolson and Schenck, 1979; Schenck et al., 1984), but has all the characteristics of the acaulosporoid morph of an *Ambispora* species (Walker et al., 1984; Spain et al., 2006).

Phylogenetic and/or morphological analyses led to a separation of Archaeosporaceae and Ambisporaceae from the acaulosporoid species (sensu stricto) of the Acaulosporaceae (Morton and Redecker, 2001; Schüssler et al., 2001; Sieverding and Oehl, 2006; Spain et al., 2006; Walker et al., 2007a; 2007b; Walker, 2008; Bills and Morton, 2015). Both families are currently important members of the Archaeosporomycetes, a fungal class, which was separated, together with Paraglomeromycetes, from the Glomeromycetes (Oehl et al., 2011a; 2011b) and comprise also the old family Geosiphonaceae (Engler and Gilg, 1924) and the recently described Polonosporaceae (Błaszkowski et al., 2021). In the following years, two remarkable species were described in the Ambisporaceae, bimorphic *Am. granatensis* (Palenzuela et al., 2011), - and *Am. reticulata* (Oehl et al., 2012), for which three-walled spores, resembling the ambisporoid-acaulosporoid morph of *Ambispora*, but neither sporiferous saccules nor mono-walled, ambisporoid-glomoid spores were recognized. Here, it is summarized that typical *Ambispora* species are bi-morphic as known for instance for *Am. fennica*, *Am. gerdemannii*, *Am. brasiliensis*, *Am. appendicula*, *Am. jimgerdemannii*, and *Am. granatensis*. They, thus form three-walled ambisporoid-acaulosporoid spores generally on a branching pedicel arising from the neck or stalk of sporiferous saccules, and mono-walled ambisporoid-glomoid spores, which typically have a subhyaline to creamy or brownish outer, evanescent layer and a persistent, but white to hyaline, structural, laminate second layer. For *Am. callosa*, we re-analysed slides with spores mounted in 1987 from the type location and did not only find again the glomoid morph – as described in Sieverding (1988), but also the acaulo-appendiculosporoid morph during the analyses for the present study in January 2026. For *Am. nicolsonii*, only the ambisporoid-acaulosporoid spore type has been known so far. For other species, such as *Am. fecundispora* only the mono-walled ambisporoid-glomoid spores have been known so far, with the subhyaline to creamy or brownish outer, evanescent layer and a persistent, but white to hyaline, structural, laminate second layer, typical and diagnostic for this morph (Spain et al., 2006, Walker et al., 2007a, Goto et al., 2008, Palenzuela et al., 2011).

Finally, for *Am. reticulata* only triple-walled spores have been described so far, but these are apparently not formed on sporiferous saccules, but on subtending hyphae, and a mono-walled ambisporoid-glomoid morph is also not known for such species.

Since a few years, separation of genera in Glomerales is performed, when the molecular dissimilarities in the partial nrDNA gene reach approximately 10% between different clades (e.g. Silva et al., 2023). In other orders these percentages might be slightly smaller such as in Entrophosporales (Silva et al., 2025) or Gigasporales (Hyde et al., 2024; Oehl et al., 2026). In the Archaeosporaceae, recently new genera were separated based on phylogenetic analyses, morphology and the percentual of dissimilarities between the different clades (Esmaeilzadeh-Salestani et al., 2025). In this family *Andinospora* and *Antiquispora* were separated from the old genera *Archaeospora* (Morton & Redecker 2001) and *Palaeospora* (Oehl et al., 2015). The objectives of the present study were to perform new molecular phylogenetic analyses on the family Ambisporaceae based on sequences from the current data sets. If major clades are obtained, the new findings would be compared with the current knowledge on morphological bases for all taxa within these clades. In case that major morphological differences between clades can be attributed to the major phylogenetic clades, new genera should be described based on these differences and molecular dissimilarities. The final aim of this study was to group these concomitant phylogenetic and morphological differences within the family Ambisporaceae.

## Materials and Methods

### Phylogenetic Analyses

To reconstruct the phylogeny, three alignments (datasets), based on partial SSU, ITS region and partial LSU nrDNA (dataset 1 - SSU+ITS+LSU), ITS region of the nrDNA (dataset 2 - ITS), and complete SSU nrDNA (dataset 3 - SSU) were generated with AM fungal sequences from species of Ambisporaceae (Supplementary Material, Spreadsheet S1). *Paraglomus brasilianum* (Spain & J. Miranda) J.B. Morton & D. Redecker, *Archaeospora trappei* (R.N. Ames & Linderman) J.B. Morton & D. Redecker, *Polonospora polonica* (Błaszk.) Błaszk., Niezgoda, B.T. Goto & Magurno and *Geosiphon pyriformis* (Kütz.) F. Wettst. were included as outgroup.

*Geosiphon pyriformis* was used as outgroup just for the dataset 1 and 3; due this species does not present ITS sequences, it was not possible to include *Ge. pyriformis* in the dataset 2. *Po. polonica* was not used as outgroup for the dataset 3, because it lacks complete SSU sequences. The sequences available for *A. granatensis* are from ITS region and complete SSU nrDNA. The ITS sequences from *Am. granatensis* were used to compose the datasets 1 and 2. The datasets were aligned in MAFFT (Katoh et al., 2019) using the default parameters. Prior to phylogenetic analyses, the model of nucleotide substitution was estimated with ModelTest-NG (Darriba et al., 2020). Maximum Likelihood - ML (1000 bootstrap) analyses were performed with Felsenstein Bootstrap Proportion (FBP) and Transfer Bootstrap Expectation (TBE), using RaxML-HPC-SSE3 and RaxML-NG, respectively (Stamatakis, 2014; Kozlov et al., 2019; Edler et al., 2020). Sequences from six of the eleven AMF species currently attributed to the Ambisporaceae were available from public data bases for us, except for *Am. brasiliensis, Am. fecundispora, Am. jimgerdemannii,* and *Am. nicolsonii*. *Ambispora reticulata* has just three fragments from partial SSU nrDNA sequences. In a Blastn analysis we observed that these sequences are related to *Endogone* spp., thus, it was not possible to use this species in our work. It was not possible to obtain sequences from the new species *Pelotaspora austrolatina*. We analyzed, by querying the Blastn on NCBI (Johnson et al., 2008), the *Ambispora* species sequences, retrieving the environmental and AMF species sequences most related to each fungus.

### Specimen analyses

Specimen of almost all AMF species currently attributed to the Ambisporaceae were available for us. Type material was available for *Am. appendicula* (OSC #41,495), *Am. brasiliensis* (HURM 78879, HURM 78880, OSC# 134501, ZT MYC 159), *Am. callosa* (GOET, OSC), *Am. fennica* (specimen kindly provided by M. Vestberg), *Am. gerdemannii* (OSC #39,476), *Am. granatensis* (ZT Myc 1626, OSC #134,712), *Am. jimgerdemannii,* (OSC #37,417), *Am. leptoticha* (OSC #40,249), *Am. reticulata* (ZT Myc 24171-24175) and the species newly described within the present study (ZT Myc 15115-15119). Especially, we also re-visited the type location of *Am. gerdemannii* in October 2005 for soil and spore collection, together with Joyce L. Spain, and guided by Prof. James M. Trappe, who already was first collector at the location ‘1 km north of Benham Falls at Fort Benham’ (Rose et al., 1979). Additionally, we re-analysed about 100 slides from the type location of *Am. callosa*, basionym *Glomus callosum* (Sieverding 1988), prepared in 1987 by Ewald Sieverding and originating from an experimental field of the agricultural school Mushweshwe, near Bukavu in the South-Kivu province in Zaire (since 1997 belonging to the Democratic Republic of Kongo, Central Africa). On several of these slides, we found not only the glomoid morph of *Am. callosa*, but also an ambisporoid-aacaulosporoid morph of an *Ambispora* (sensu lato) species. In 1987, this ambisporoid-aacaulosporoid morph had been miss-identified as *Acaulospora appendicula*. Selected slides showing the later morph were deposited at Z+ZT (the common mycological herbarium of the University and ETH of Zurich, Switzerland) with the accession number ZT Myc 94416, while newly preprared slides from the type location of *Am. gerdemannii* were deposited under the accession (ZT Myc 94417). The specimens for the present study were either observed directly in compound microscopes in the mycological herbaria (e.g. OSC) or in living culture collections (e.g. INVAM), after loan, or from our own institutional, or private collections (E. Sieverding; F. Oehl) in Basel, Zürich, Granada or Temuco (Chile).

For the description of a new species, or morphological analyses on other, already described *Ambispora* spp., also fresh spores were extracted, either directly from field soil samples or from trap cultures, maintained within the institutional greenhouses in Basel and Zürich (Switzerland), Granada (Spain), Tarapoto (Peru) and Recife (Brazil), by wet-sieving and sucrose centrifugation (Sieverding, 1991). Such spores were immediately mounted on slides. When described by us, about 100 spores were examined per species. Spore formation characteristics and morphology of spores were observed on specimens mounted in polyvinyl alcohol-lactic acid-glycerol (PVLG) (Koske and Tessier, 1983), in a mixture of PVLG and Melzer’s reagent (Brundrett et al., 1994) and in water (Spain, 1990). Specimens mounted in PVLG and the mixture of PVLG and Melzer’s reagent were deposited at Z+ZT and HURM, the mycological herbarium of the UFPE in Recife. All spore observations and information on spore characteristics are based on spores extracted from soil, trap cultures, or single, or multiple spore-derived pure cultures. No information is provided from in-vitro-cultured materials. Spore wall terminology follows the nomenclature of Spain (2003), Spain et al. (2006), Goto et al. (2008) and Palenzuela et al. (2011). Analyses of the spore walls, germination structures and mycorrhizal structures were performed using compound microscopes at 100-1000 x magnifications. For this paper, all original species descriptions and available species emendations were also considered.

### Soil sampling for the newly described species

Soil samples were taken with a shovel in a Southern Chilean temperate grassland and a South Chilean deciduous forest (Castillo et al., 2006), in two conventionally managed agricultural soils from the rhizosphere of *Triticum aestivum* (September 2011), respectively, and in a conventionally managed South Brazilian vineyard the rhizosphere of *Vitis vinifera* (September 2004). The grassland and the forest site were located in the Experimental Station San Pablo de Tregua of the University Austral de Chile (Valdivia, Chile, 39°30-39°38’S and 72°02-72°09’W) at an elevation between 550 and 1600 m asl (Castillo et al., 2006). The two wheat fields were located in Curacautín (38°51’S, 71°83’W at 605 m asl) and Gorbea (39°12’ S, 72°59’ W at 179 m), two municipalities in the Araucanía region of Southern Chile, and the vineyard in Caxias do Sul, Rio Grande do Sul, at about 825 m asl (29° 16’ 33”S and 51° 07’ 06”W; Silveira, 2006). The soil pH (H2O) was 5.4 in the grassland and the deciduous forest, 5.9 and 4.8 in the wheat fields, and 6.6 in the vineyard. Organic carbon was 88.7 mg kg^-1^ in the grassland, 144.4 mg kg^-1^ in the deciduous forest, 136.0 mg kg^-1^ and 95.0 mg kg^-1^ in the wheat fields, and 44.0 mg kg^-1^ in the vineyard. Available P was 2.9 and 3.6 mg kg^-1^ in the grassland and the deciduous forest, respectively, 47.8 mg kg^-1^ and 31.9 mg kg^-1^ in the wheat fields, while it was >100.0 mg kg^-1^ in the vineyard. All Chilean soils were Andosols, while the soil type of the vineyard in Southern Brazil was a Leptosol.

## Results

### Molecular phylogeny

Currently Ambisporaceae is monogeneric with the single genus *Ambispora*. According to our phylogenetic analyses, Ambisporaceae present three different clades well supported (Figs. 1, 2, and 3). The clade ‘A’ is represented by *Am. fennica*, and a sequence attributed to *Am. gerdemannii*. The clade ‘B’ is composed by sequences from *Am. callosa*, *Am. leptoticha*, *Am. appendicula*, and two sequences attributed to *Am. gerdemannii* (isolate AU215) in the NCBI (Fig. 2). The clade ‘C’ has, so far, sequences from *Am. granatensis*. In our opinion the clades ‘B’ and ‘C’ represent two new genera.

**Fig. 1:**
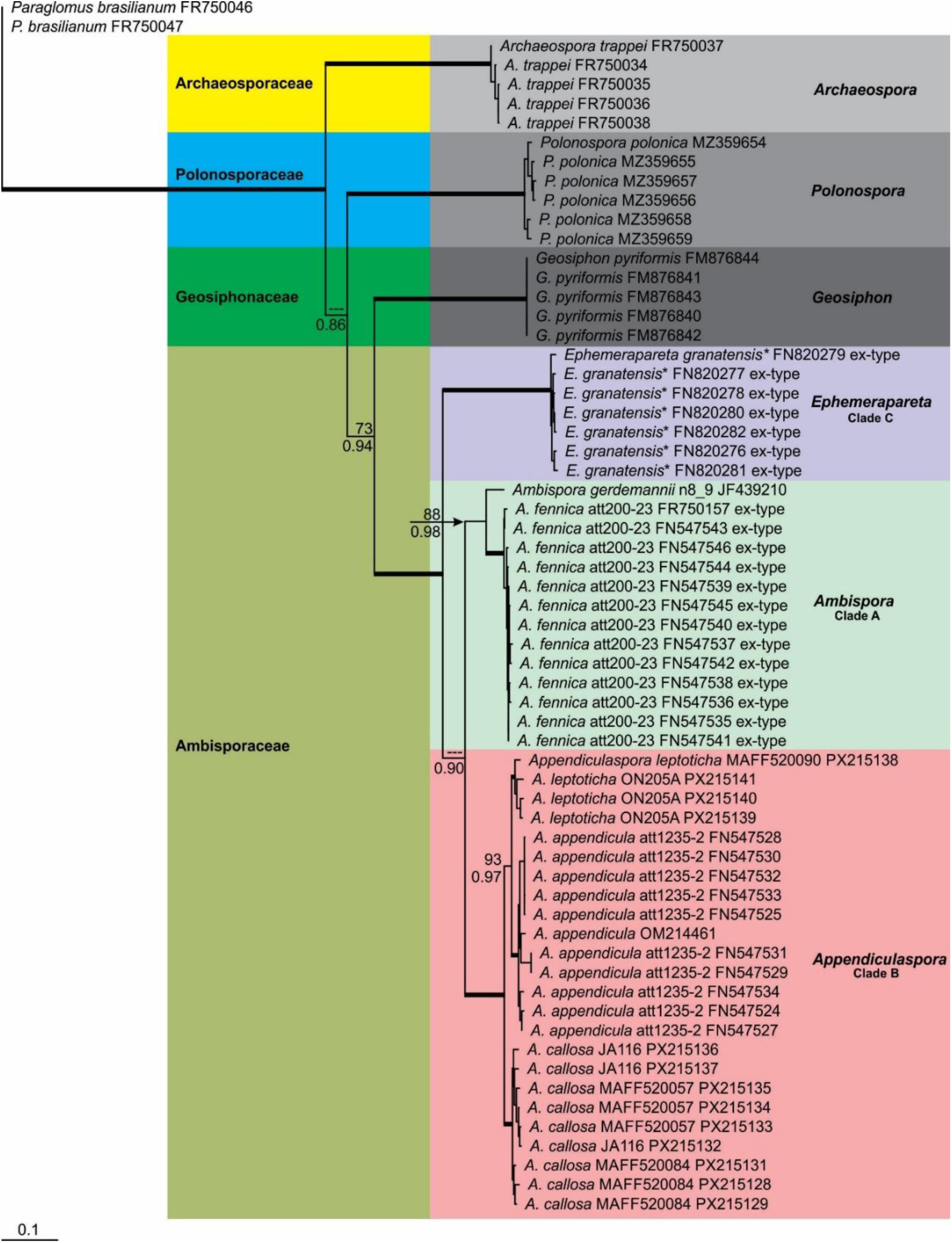
Phylogenetic tree obtained by analysis from partial SSU, ITS region, and partial LSU nrDNA sequences of Ambisporaceae spp. Sequences are labelled with their database accession numbers. Support values (from top) are from ML (maximum likelihood analysis) using RaxML-HPC-SSE3 with FBP (Felsenstein Bootstrap Proportion) and RaxML-NG with TBE (Transfer Bootstrap Expectation), respectively. Only support values of at least 70% are shown. Thick branches represent clades with more than 95% of support in all analyses. The tree was rooted by *Paraglomus brasilianum*, *Archaesopora trappei*, *Polonospora polonica* and *Geosiphon pyriformis*. Sequences with only the ITS region are indicated by *. The nucleotide substitution model used was GTR+I+G for both analyses.

**Fig. 2:**
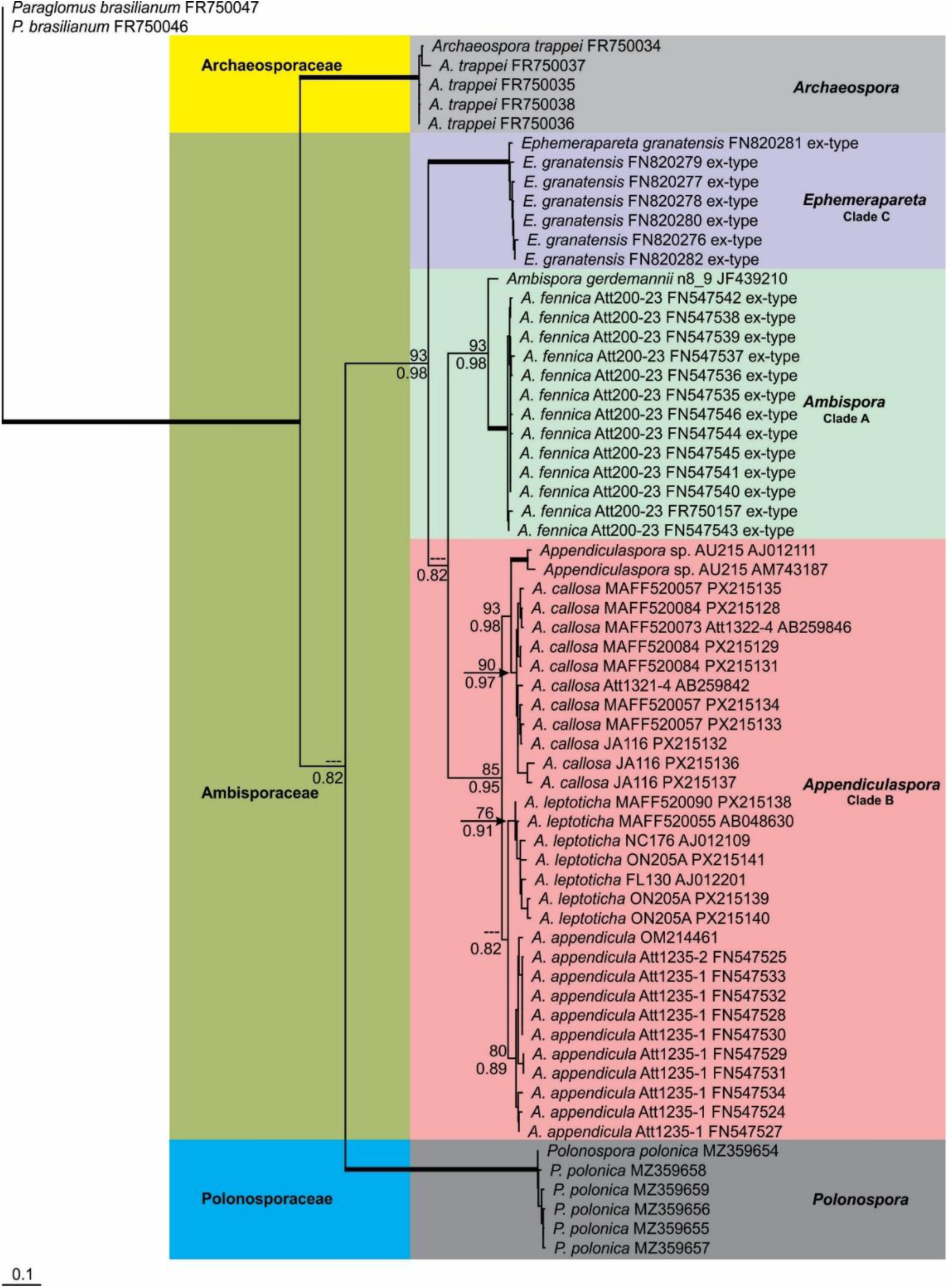
Phylogenetic tree obtained by analysis from ITS nrDNA sequences of Ambisporaceae spp. Sequences are labelled with their database accession numbers. Support values (from top) are from ML (maximum likelihood analysis) using RaxML-HPC-SSE3 with FBP (Felsenstein Bootstrap Proportion) and RaxML-NG with TBE (Transfer Bootstrap Expectation), respectively. Only support values of at least 70% are shown. Thick branches represent clades with more than 95% of support in all analyses. The tree was rooted by *Paraglomus brasilianum*, *Archaeospora trappei*, and *Polonospora polonica*. The nucleotide substitution model used was GTR+G for RaxML-HPC-SSE3 and HKY+G for RaxML-NG analyses.

**Fig. 3:**
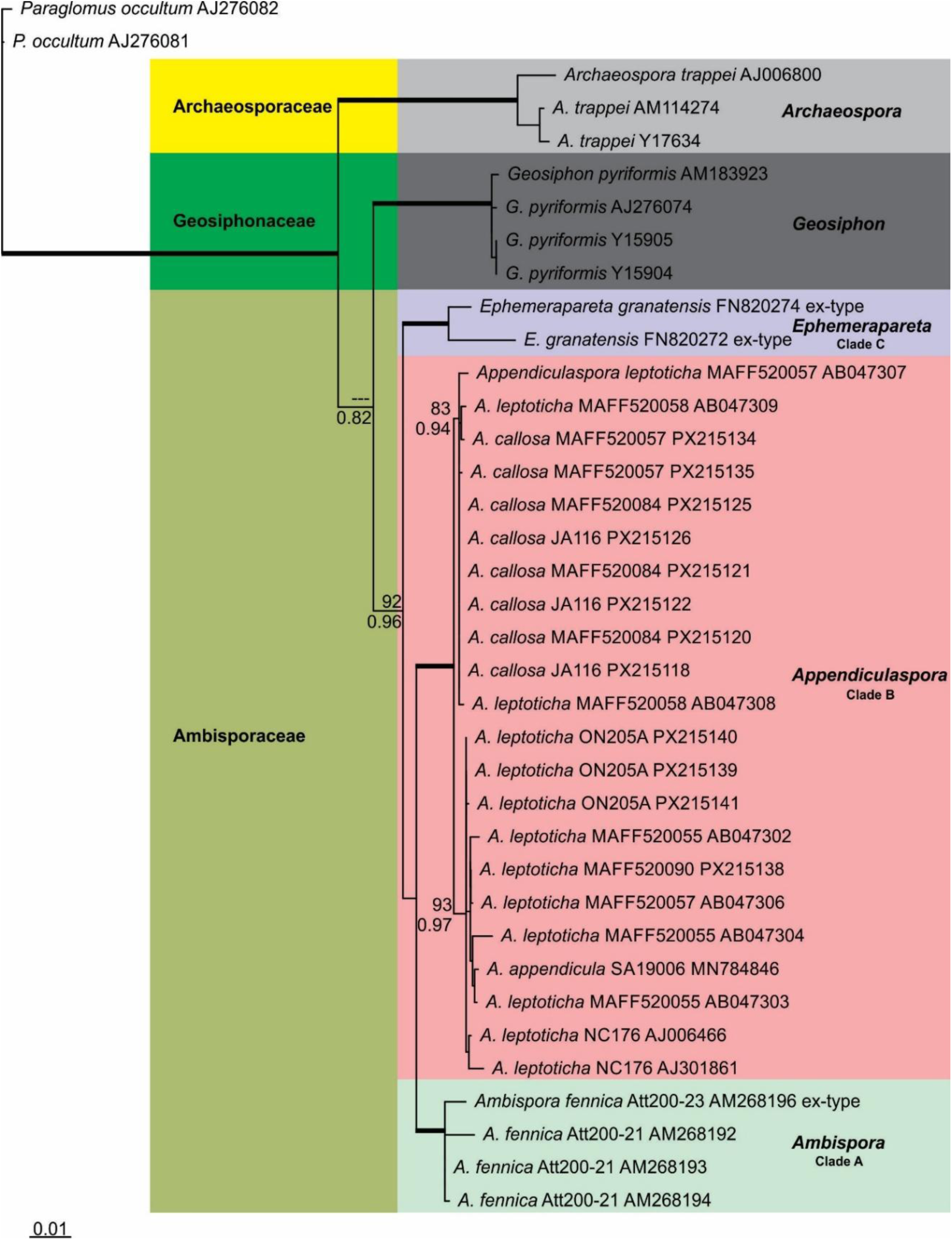
Phylogenetic tree obtained by analysis from SSU nrDNA sequences of Ambisporaceae spp. Sequences are labelled with their database accession numbers. Support values (from top) are from ML (maximum likelihood analysis) using RaxML-HPC-SSE3 with FBP (Felsenstein Bootstrap Proportion) and RaxML-NG with TBE (Transfer Bootstrap Expectation), respectively. Only support values of at least 70% are shown. Thick branches represent clades with more than 95% of support in all analyses. The tree was rooted by *Paraglomus brasilianum*, *Archaeospora trappei*, and *Geosiphon pyriformis*. The nucleotide substitution model used was GTR+I+G for RaxML-HPC-SSE3 and TIM+I+G for RaxML-NG analyses.

In Ambisporaceae sequences from *Am. appendicula*, *Am. callosa*, *Am. leptoticha*, *Am. gerdemannii* and *Am. fennica* are available with the entire fragment (partial SSU + ITS region + partial LSU nrDNA), analysed in the dataset 1. According to Maximum Identity (MI), in the Blastn analyses (considering the entire fragment - SSU-ITS-LSU), the species sequences closest to the clade ‘A’ (*Am. fennica*; *Am. gerdemannii*) are from *Am. callosa* - MAFF520057 (89% MI), *Am. appendicula* - att1235-2 (88.9% MI), and *Am. leptoticha* - MAFF520090 (88.2% MI). Querying the ITS sequences from *Am. granatensis* (clade ‘C’, new genus *Ephemerapareta*) with the Blastn, the nearest sequences (85% MI) are from *Am. fennica*, but with an e-value of 2e-117, indicating that the ITS sequences from *Am. granatensis* are not close to any other species in Ambisporaceae. Thus, the nearest species from clade ‘B’ (new genus *Appendiculospora*), in relation to clade ‘A’ (*Ambispora* sensu stricto), has 89% MI, and no species from the clade ‘A’ or ‘B’ is close to *Am. granatensis* (clade ‘C’).

Considering the entire fragment (partial SSU + ITS region + partial LSU nrDNA), we found three environmental sequences related to *Ambispora* (sensu stricto) with ≥ 95% MI. Two of these environmental sequences were obtained from roots of *Littorella uniflora* (LS479906, LS479907) in Norway (Sudová et al., 2021), and one from roots of *Sequoiadendron giganteum* (HQ895806) in USA (Fahey et al., 2012). In relation to the new genus 1 (clade B - *Appendiculaspora*), we found six environmental sequences with ≥ 95% MI. Four of these sequences were obtained from roots of *Solanum tuberosum* (HF970299, HF970300, HF970301, HF970302) in Peru (Senés-Guerrero et al., 2013), and two from rhizosphere soil of *Sinocalycanthus chinensis* (MH469316, MH469376) in China (not published). No environmental ITS sequences were found related to *Am. granatensis* (clade C, new genus 2 - *Ephemerapareta*).

### Morphological analyses

Ambisporaceae species have generally been known to be bi-morphic (see Tab. 1), as spores can be formed i) ambisporoid-acaulosporoid, i.e. terminally on the appendix of a branching neck of a sporiferous saccule, which is also formed terminally, and ii) terminally on subtending hyphae (i.e. typical ambisporoid-glomoid morph). The phylogenetic analyses request for advanced morphological separations. These can primarily be based on spore wall composition of the triple-walled ambisporoid-acaulosporoid morph, although the outer wall is semi-permanent to evanescent in all *Ambispora* spores of this morph: I) *Am. fennica*, *Am. brasiliensis*, *Am. gerdemannii* and *Am. nicolsonii* have a smooth, permanent central spore wall (*Am. fennica* clade, ‘A’, Figs 4-9), while ii) the central wall of *Am. appendicula*, *A. leptoticha*, *Am. jimgerdemannii* and also *Am. callosa* is alveolate (*Am. appendicula* clade, ‘B’, Figs 10-19), and iii) the central wall of *Am. granatensis* is smooth, but easily degraded, thus rather short-lived and not permanent, but also evanescent (*Am. granatensis* clade, ‘C’, Figs 20-22). Species of the *Am. fennica* (clade ‘A’) represent the genus *Ambispora* (Tab. 1), based on the type species *Am. fennica*, while species of the *Am. appendicula* (clade ‘B’) represent the new genus *Appendiculaspora* (Tab. 1), and the mono-specific *Am. granatensis* (clade ‘C’) represents the new genus *Ephemerapareta* (Tab. 1). However, species for which hitherto only an ambisporoid-glomoid morph has been known, can currently not be attributed to one of these clades, based on this distinction of the acaulosporoid morph, and thus can only be identified readily by phylogenetic analyses on the genus level. Such species currently must remain in the type genus *Ambispora* of the revised Ambisporaceae.

**Tab. 1:**
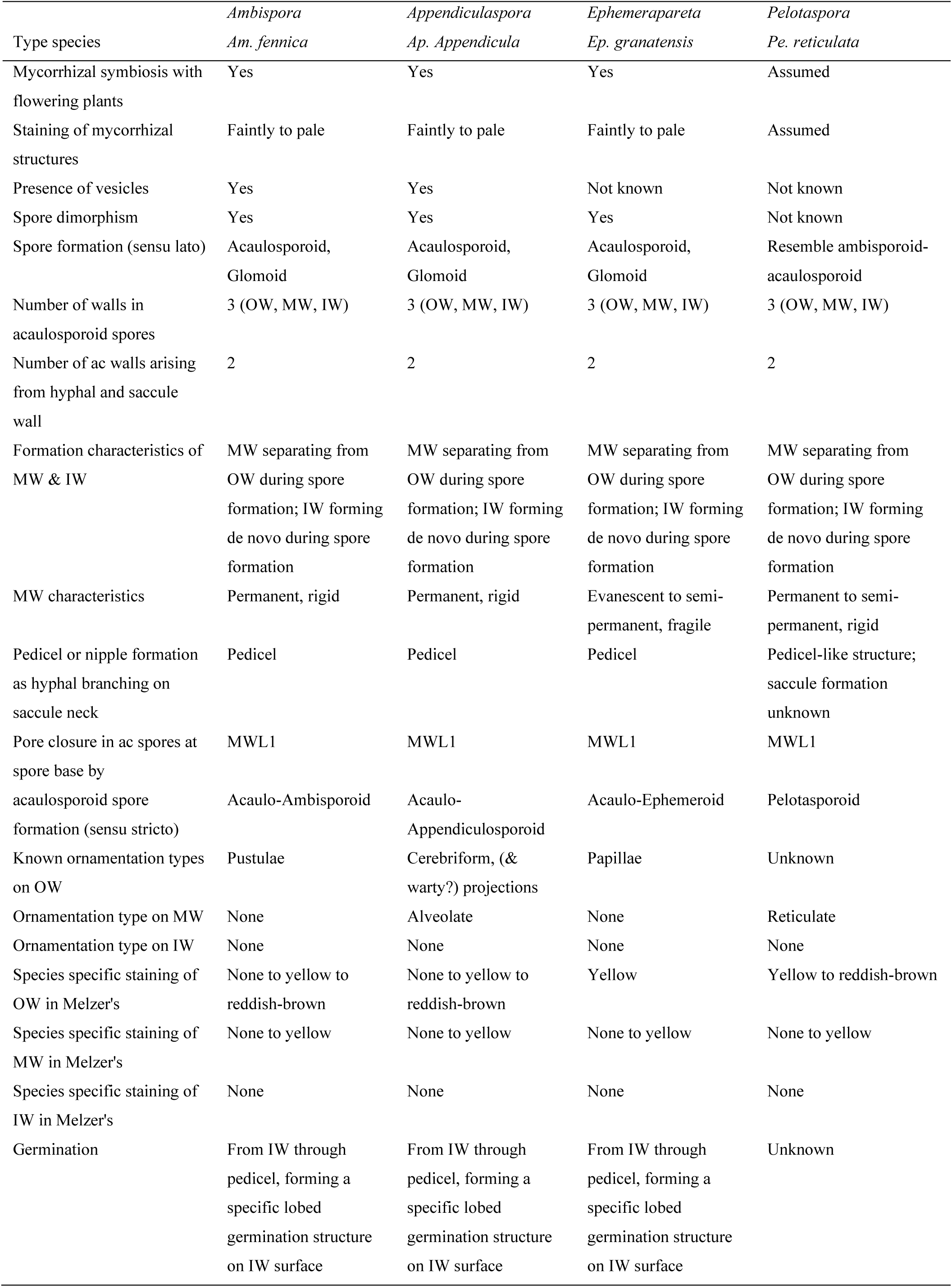
Major morphological characteristics separating genera for the triple-walled morph of the Ambisporaceae.

One additional morph, with triple-walled spores, but apparently formed on subtending hyphae, has a diagnostic reticulate, football-like middle wall (Figs 23-35). It is represented by *Am. reticulata* (Figs 23-26), for which so far all molecular analyses on field-soil sampled or trap cultures have failed. Thus, a new description of this taxon on the genus level can solely be based on morphological analyses. A similar species (Figs 27-35) had been found from Southern Chile and Southern Brazil, respectively, but molecular identification analyses also failed so far for this species. Due to their congruent spore morphology, well different from those of *Ambispora* (sensu lato) clades ‘A’-‘C’ (Tab. 1), here it was decided to attribute both species to a new genus within the family, rather than to attribute both erroneously to one of the other three genera, to which they cannot anymore be attributed due to the advanced morphological analyses presented here.

**Figs. 4-9:**
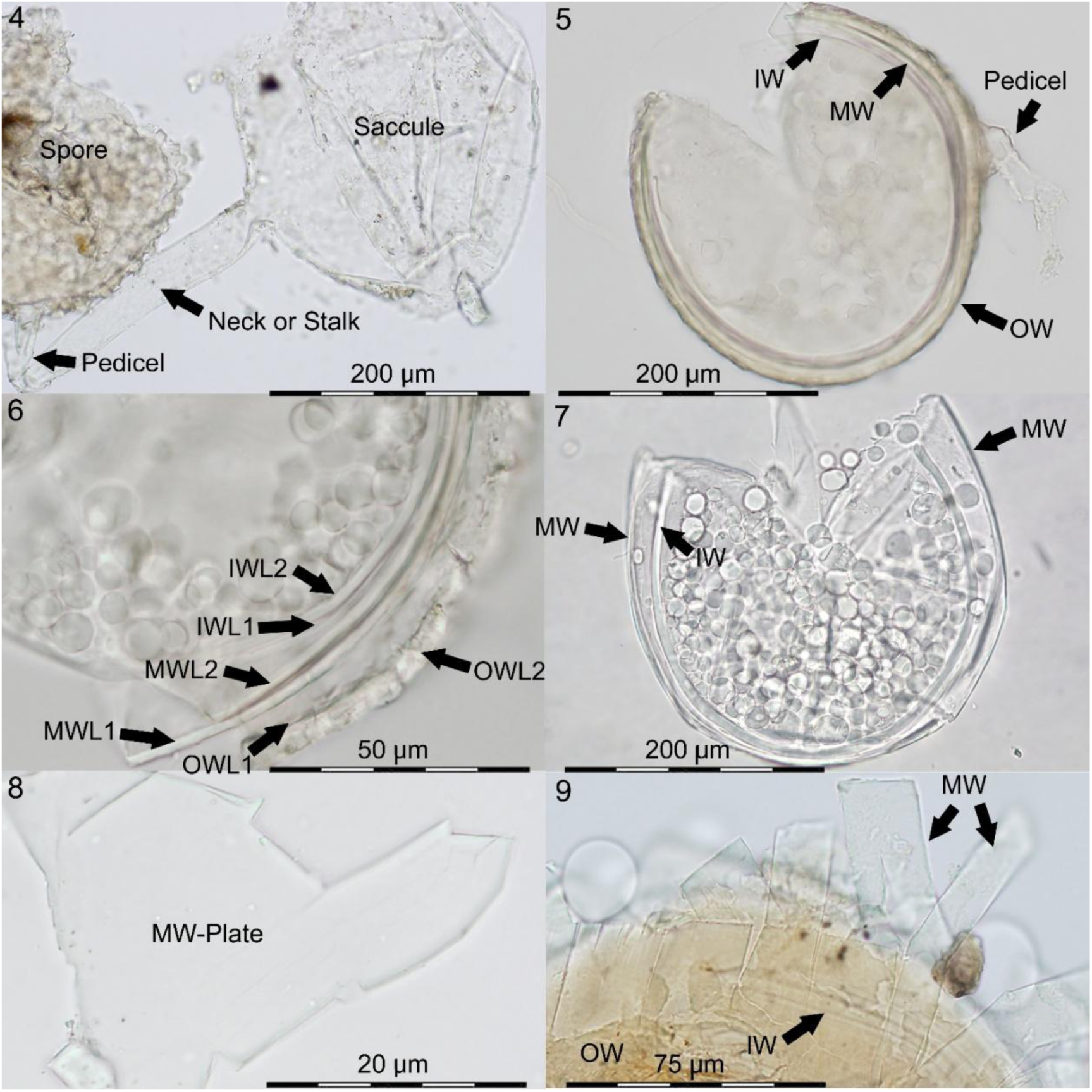
Acaulo-ambisporoid spores and spore characteristics of *Ambispora* spp.: Figs. 4-8: *Ambispora fennica*. Fig. 4. Spore, formed on a hyaline pedicel, branching from the neck, or stalk of a sporiferous saccule. Fig. 5. Crushed spore with OW, MW & IW, showing the pedicel, formed by hyaline MWL1 and a plate, splitted from the crushed MW at the top. Fig. 6. Crushed spore segment with typically fissured OW, bi-layered MW (MWL1 & MWL2) and bi-to-triple layered IW. Fig. 7. OW completely lost from the spore, Fig. 8. M-plate, splitted in irregular way, but showing several sharp edges and corners. Fig. 9. *Ambispora gerdemannii*. Plate-like splitting on crushed MW. Here, the plates splitted into regular rectangles.

**Figs. 10-17:**
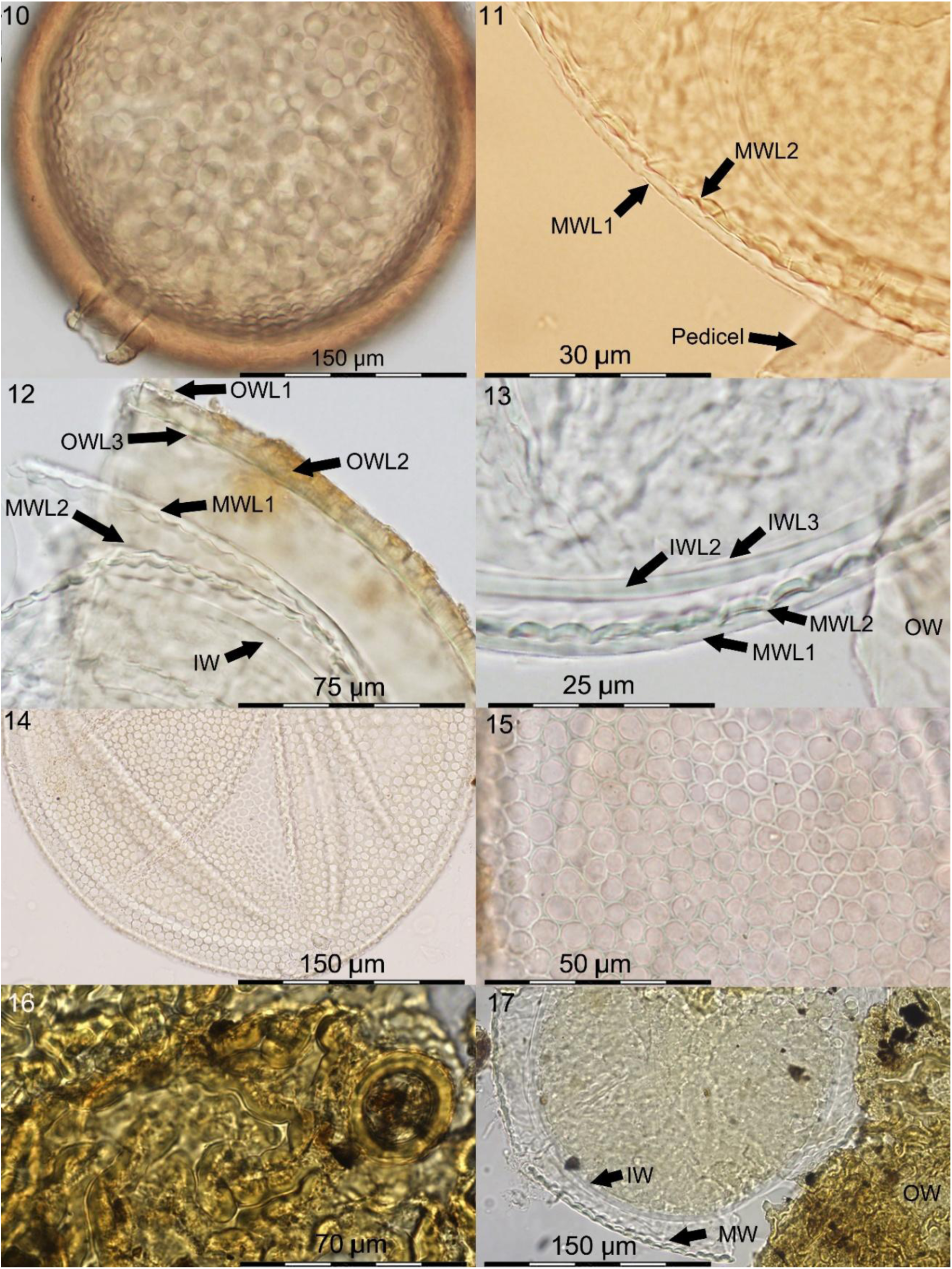
Acaulo-appendiculosporoid spores and spore characteristics of twp *Appendiculaspora* spp.: Figs. 10-15: *Appendiculaspora appendicula*. Fig. 10. Spore with pigmented OW (surface), alveolate MW beneath and pedicel formed by both walls. Fig. 11. Alveolate MW consisting of two layers. MWL1 does not stain in Melzer’s, but MWL2 stains yellowish. Fig. 12. Crushed spore with smooth, but fissured OW, alveolate MW and smooth IW. Fig. 13. Alveolate MWL1 & MWL2, and bi-to-triple-layered IW. Figs. 14-15. Alveolate MW in planar view. Figs. 16-17: *Appendiculaspora jimgerdemannii*. Fig. 16. Labyrinthiforme/cerebriforme surface ornamentation of OW. A round collar of OW is well visible on the right, thus pedicel has broken away at the collar. Fig. 17. Ornamented OW, alveolate MW and smooth IW.

**Figs. 18-24:**
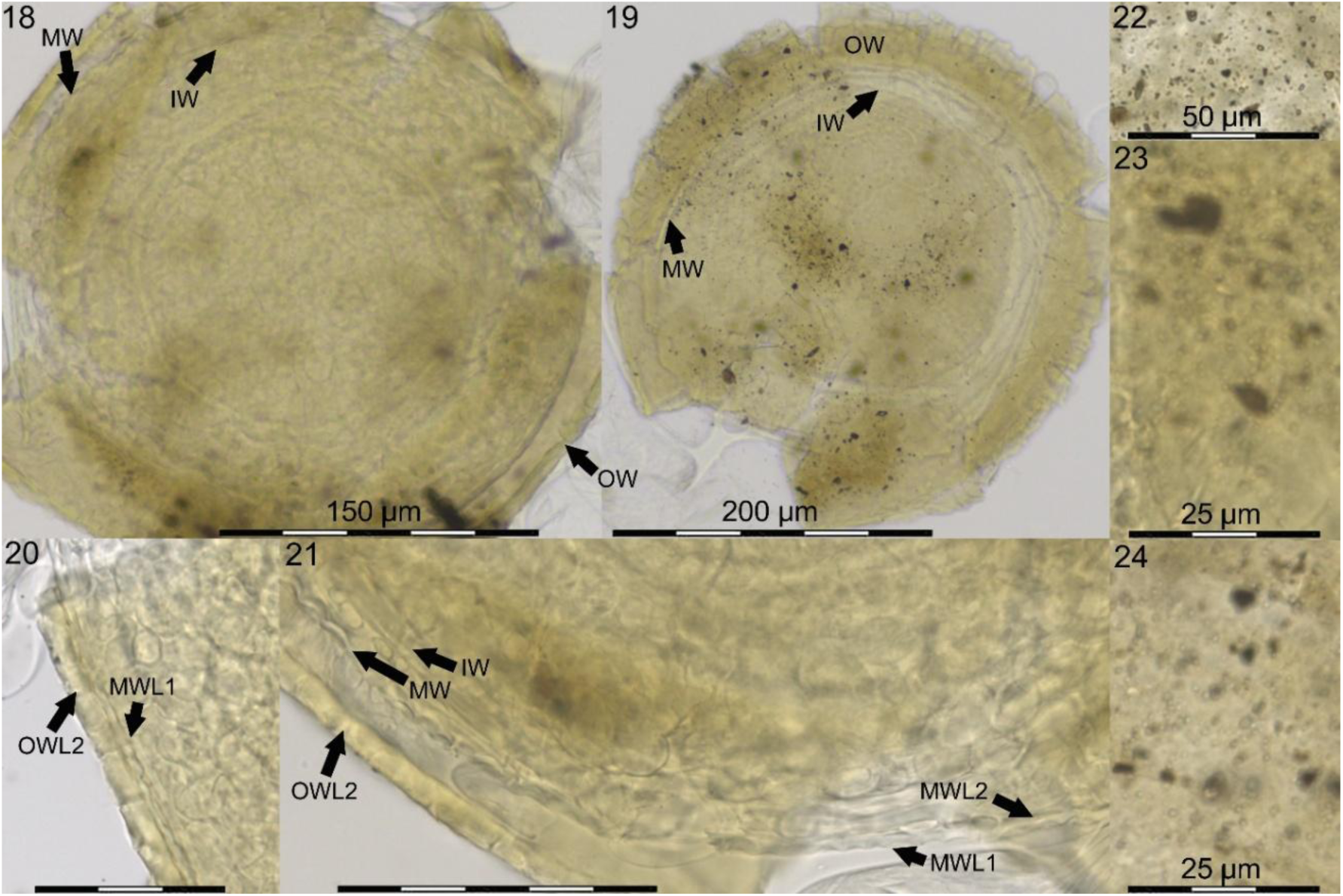
*Appendiculaspora callosa.* Figs 18-21. Crushed spores; Outer wall (OW) subhyaline to light creamy. OWL2 finely laminate, first rigid, but not permanent, as aged (Fig. 19) showing multiple fissures, degraded. MW (with MWL1 & MWL2) alveolate; IW smooth. Figs 22-24. Minute (spiny) warts on the OW spore surface in planar view (about 0.5 μm in diam).

**Figs. 25-27:**
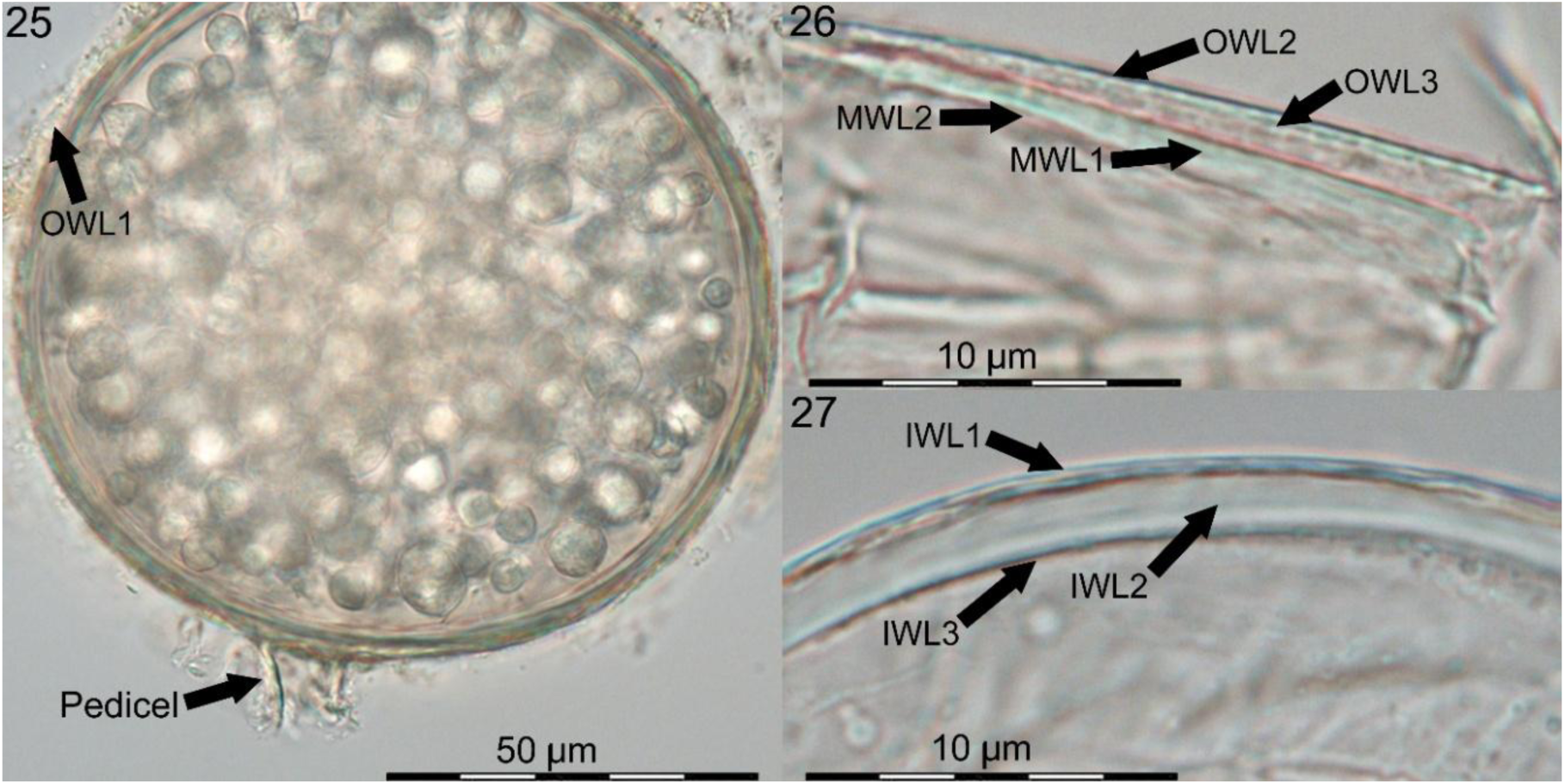
Acaulo-ephemeroid spores and spore characteristics of *Ephemerapareta granatensis*. Fig. 25. Spore with pedicel attached. Fig. 26. Short-lived, evanescent OW (rather thin, with OWL1 & OWL2) and evanescent to semi-permanent MW (MWL1 & MWL2). Fig. 27. Permanent IW (IWL1-3).

**Fig. 28-31:**
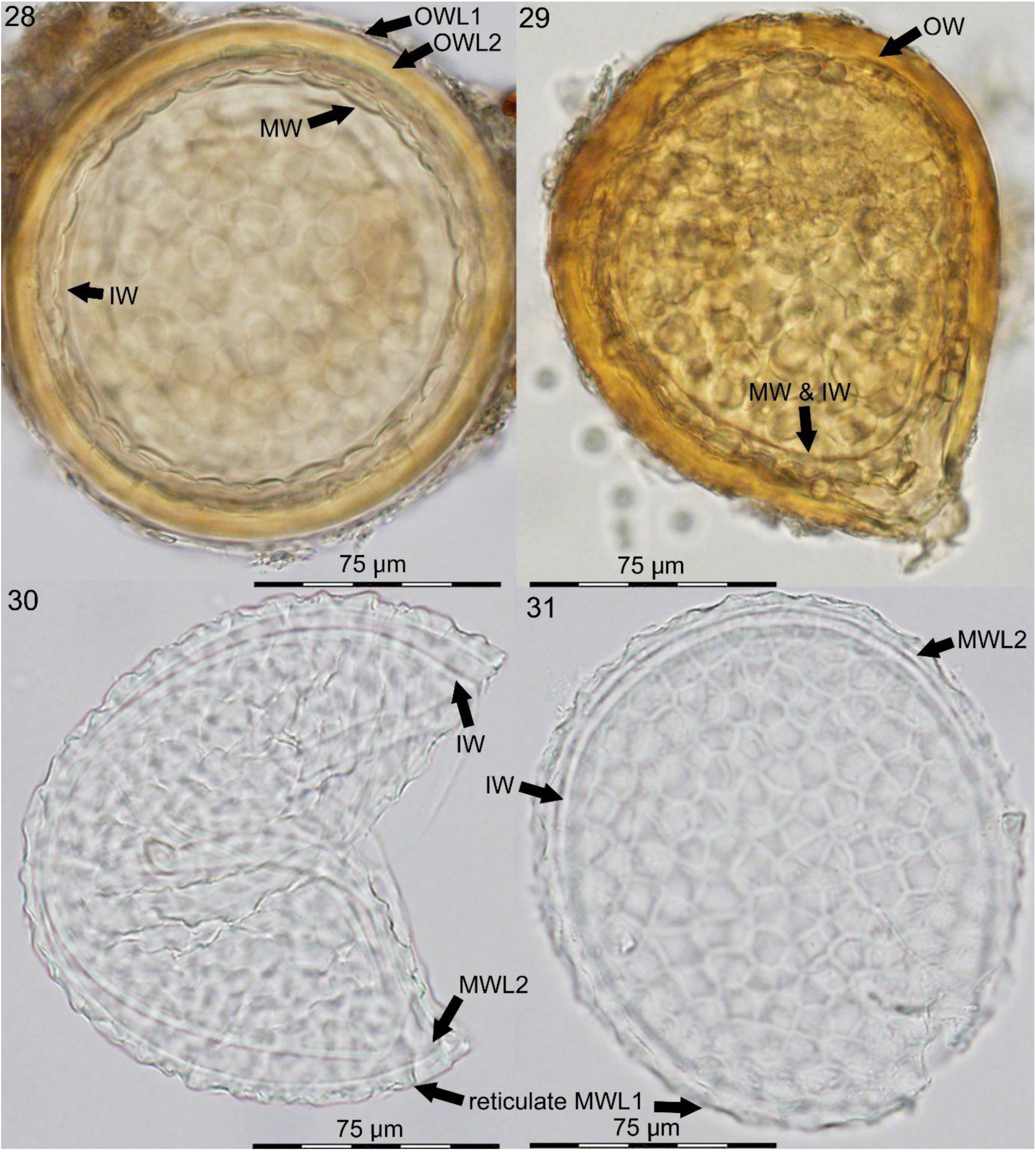
*Pelotaspora reticulata*. Figs. 28-29. Spores with pigmented, smooth OW, with subhyaline OWL1 & brown OWL2, reticulate MW, and smooth IW, and a pedicel-like structure at the spore base (Fig. 23). Figs 30-31. Reticulate MWL1, smooth MWL2 and smooth IW. The reticulum consists of (4-)5-6(−8) sided, large pits (15-30 x 12-25 µm) that are framed by 3-8 µm broad ridges giving to the spore a football-like appearance.

**Figs. 32-40:**
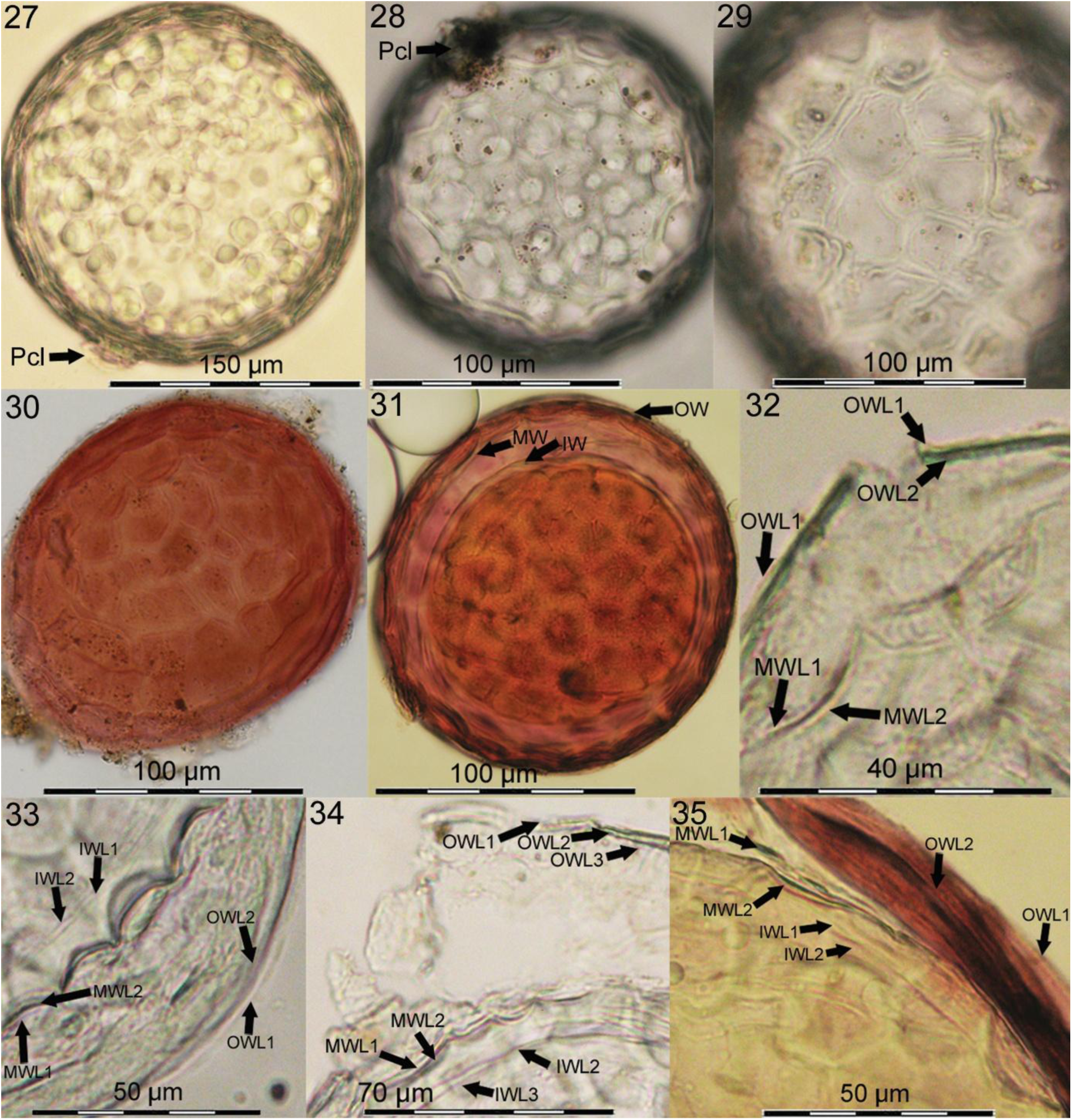
*Pelotaspora austrolatina*. Figs. 32-34. Spores with a pedicel-like (Pcl) subtending hyphae and reticulate appearance deriving from the reticulum on the MW surface. OW surface is smooth. Figs. 35-36. Spores in Melzer’s reagent: OW staining purple. IW yellowish. Fig. 37-39. Crushed spore in PVLG, with smooth, evanescent triple layered OW, reticulate bi-layered MW, and smooth bi-to-triple layered IW. Fig. 40. Crushed spore in PVLG+Melzer’s: OWL2 staining purple and MWL2 staining yellowish.

### Taxonomy

**Ambisporaceae** C. Walker, Vestberg & A. Schüßler, Mycol. Res. 111 (2): 143 (2007), emended here by Oehl. G.A. Silva, Palenz. & Sieverd.

MycoBank MB 510208

Emended description: Sporocarp formation unknown; species generally bimorphic with mycorrhizal associations producing both three-walled acaulosporoid and generally mono-walled glomoid morphs on extra- or intraradical hyphae, but acaulosporoid or glomoid morph is not (yet) known for all species. *Acaulosporoid spores* on short hyphal appendix generally arising laterally from the hyphal neck of a terminal sporiferous saccule; with outer, middle and inner wall; germinating from the innermost wall with a germ tube emerging through the appendix attachment or penetrating the outer walls; a germination structure possibly formed between inner and middle wall. *Glomoid spores* formed singly or in loose clusters on extra- or intraradical hyphae, with a hyaline to subhyaline to creamy or brown outer layer and a hyaline to white structural layer beneath, rarely with triple walls as known for the acaulosporoid morph; germinating through the subtending hypha. Intraradical mycorrhizal hyphae, arbuscles and vesicles stain pale blue with trypan blue.

Type genus: ***Ambispora*** C. Walker, Vestberg & A. Schüßler, Mycol. Res. 111 (2): 147 (2007) MycoBank MB 510209

Other genera: *Appendiculaspora*, *Ephemerapareta*, *Pelotaspora*

***Ambispora*** C. Walker, Vestberg & A. Schüßler, Mycol. Res. 111 (2): 147 (2007), emended here by Oehl, Sieverd., Palenz. & G.A. Silva, Figs 4-9

MycoBank MB 510209

Emended description: Sporocarp formation unknown. Acaulosporoid spores formed singly on short hyphal appendix generally arising laterally from the hyphal neck of a terminal sporiferous saccule. Glomoid spores formed singly or in loose clusters on extra- or intraradical hyphae. Sporiferous saccules formed terminally on hyphae. The saccule wall is bi- to triple layered, hyaline to subhyaline. Acaulosporoid spores have three walls: outer, middle, inner wall: OW, MW, IW; OW with two to three wall layers (OWL1-3) that may sometimes be lost totally, as they generally are evanescent to semi-permanent. MW is bi-layered (MWL1-2) and permanent. IW is three-layered (IWL1-3). OWL1 is evanescent, only present on young spores; OWL2 is semi-permanent to evanescent, easily seen, often showing a crazed surface with age; OWL3, also semi-permanent to evanescent, thin-layered, tightly pressed to OWL2. In young spores, OW and MWL1 continuous with the appendix wall and mycelial hyphae; MWL1 can produce several septa in the appendix. In mature spores where the appendix is broken off, a septum formed by MWL1 and MWL2 closes the pore at the appendix base. MWL1 and MWL2 with smooth surfaces, but often with fracturing character (i.e. ‘plate-like splitting’, irregular to rectangular splitting) in crushed spores. IWL2 finely laminate, structural layer, while IWL1 and IWL3 thin, unite, flexible and rather difficult to detect. Germination of acaulosporoid spores of *Ambispora* spp. has not yet been observed to our knowledge. Glomoid spores hyaline to subhyaline, globose to subglobose,, with one spore wall (SW), consisting of a mucilaginous, hyaline to creamy to brown outer layer SWL1, about 1.0-2,5 μm thick, frequently with adhering debris on the outer surface; inner wall layer SWL2 several μm thick, laminated, hyaline to white. Subtending hypha cylindric to funnel shaped, 5-30 μm diam at the spore base, bi-layered. Type species: *Ambispora fennica* C. Walker, Vestberg & A. Schüßler, Mycol. Res. 111 (2): 148 (2007), Figs 4-8

MycoBank MB 510210

Other species: *Ambispora brasiliensis*, *Ambispora fecundispora, Ambispora gerdemannii*, *Ambispora nicolsonii Ambispora brasiliensis* B.T. Goto, L.C. Maia & Oehl, Mycotaxon 105: 13 (2008)

MycoBank MB 511612

*Ambispora fecundispora* (N.C. Schenck & G.S. Sm.) C. Walker, Mycol. Res. 112 (3): 298 (2008)

MycoBank MB 511419

Basionym: *Glomus fecundisporum* N.C. Schenck & G.S. Sm., Mycologia 74 (1): 81 (1982)

MycoBank MB 110703

Synonym: *Appendicispora fecundispora* (N.C. Schenck & G.S. Sm.) C. Walker, Vestberg & A. Schüßler, Mycol. Res. 111 (3): 254 (2007)

MycoBank MB 510502

*Ambispora gerdemannii* (S.L. Rose, B.A. Daniels & Trappe) C. Walker, Vestberg & A. Schüßler, Mycol. Res. 111 (2): 148 (2007), Fig. 9

MycoBank MB 510211

Basionym: *Glomus gerdemannii* S.L. Rose, B.A. Daniels & Trappe, Mycotaxon 8(1): 297 (1979)

MycoBank MB 314598

Synonym: *Appendicispora gerdemannii* (S.L. Rose, B.A. Daniels & Trappe) Spain, Oehl & Sieverd., Mycotaxon 97: 174 (2006)

MycoBank MB 510321

*Ambispora nicolsonii* (C. Walker, L.E. Reed & F.E. Sanders) Oehl, G.A. Silva, B.T. Goto & Sieverd., Mycotaxon 117: 431 (2011)

MycoBank MB 561663

Basionym: *Acaulospora nicolsonii* C. Walker, L.E. Reed & F.E. Sanders, Trans. Br. mycol. Soc. 83(2): 360 (1984)

MycoBank MB 105905

***Appendiculaspora*** gen. nov. Sieverd., Oehl, Palenz. & G.A. Silva, Figs 10-24 MycoBank MB 862213

Description: Sporocarps unknown. Acaulosporoid spores formed singly upon a short appendix that arises laterally on the tapering hyphal neck of a sporiferous saccule, and glomoid spores formed terminally on hyphae, occurring singly or in loose clusters on extra- or intraradical hyphae. Acaulosporoid sporiferous saccules also formed on mycelial, extra- or rarely intraradical hyphae. Both acaulosporoid and glomoid spores may be found on the same mycelium and both can function as colonization propagules. Sporiferous saccule wall consists of one wall with 2-3 wall layers; prior to appendix and spore formation; a septum forms in the tapering hypha distal to the saccule where the hyphal appendix is formed. Spores are white-opaque to dull yellow-cream to orange-tan or creamy to light brown. Three spore walls are present: outer, middle, and inner (OW, MW, IW). OW comprises three layers in young, fully developed spores. OWL1 is hyaline, often lost or degrades during the early stages of spore formation. OWL2 is initially white, becoming yellow to brown, firm and difficult to break on young spores, but with age becoming less rigid, somewhat roughened, with an irregular crazed pattern of fine cracks. OWL3 usually is hard to observe as it is very thin and often tightly adherent to OWL2. When the appendix is broken off an open pore is formed in the outer wall that is 20-50 μm wide. MW comprises two layers: the outer layer MWL1 is continuous with the thick, hyaline layer of the appendix and second layer of the connected hypha and sporiferous saccule, and hyaline, and with a convex, alveolate reticulum. These undulations are about 5-12 μm wide and 2-6 μm deep. MWL2 is hyaline, tightly adherent to MWL1 and thus showing a similar alveolate structure with concave hemispherical depressions on the outer surface that fit into the convex protuberances at the inner surface of MWL1. MWL2 may close the pore of the appendix. MWL1 and MWL2 separate when pressure ruptures the spore. MWL2 may stain yellow in Melzer’s reagent. IW is hyaline, smooth, forms only after differentiation and complete formation of the outer and middle walls. A very thin outer layer IWL1 and a thin inner layer IWL3 appear to adhere to the finely laminated middle layer IWL2 in water mounted specimens, but neither IWL1 nor IWL3 are usually detected in specimens mounted in PVLG. The pedicel (appendix) arises laterally from the hyphal neck of a sporiferous saccule, is often persistent, and resembles a subtending hypha, 25-120 μm long, 20-50 μm wide, cylindric or funnel-shaped tapering to 10-25 μm at the distal end from the spore. The appendix pore is closed by a septum arising from MWL1 at a short distance (0-10 μm) from the spore base. Acaulosporoid spores germinate with a single or branched germ tube, 6-12 μm in diameter, that emerges from the inner wall and generally exits through the pore of the appendix. A distinctive germination structure was also observed to form between middle and inner walls in a spore mounted in water from which a germ tube emerged. Single or loose clusters of swollen hyphal tips often form on the germ hyphae at a short distance (100-200 μm) from the base of the acaulosporoid spore. Glomoid spores are hyaline to subhyaline, with one spore wall (SW) consisting of a mucilaginous outer SWL1, regularly about 1.5-2.5 μm thick, frequently with adhering debris on the outer surface, and an about 2-15 μm thick, laminated inner wall layer SWL2. Subtending hypha cylindric to slightly funnel shaped, 7-35 diam at the spore base tapering to 5-15 μm within 7-120 μm distance. The pore usually is open, 5-25 μm wide, but sometimes a thin septum deriving from the inner hyphal wall layer is seen at a short distance from the spore base. Glomoid spores germinate through the subtending hypha. Forming vesicular-arbuscular mycorrhiza, but arbuscules, vesicles, and intraradical hyphae stain pale blue with trypan blue. Etymology: Appendicula-(appendix) and -spora (spore) referring to the type of spore formation on a pedicel (appendix)

Type species: *Appendiculaspora appendicula* (Spain, Sieverd. & N.C. Schenck) Sieverd., G.A. Silva & Oehl, comb. nov., Figs 10-15

MycoBank MB 862214

Basionym: *Acaulospora appendicula* Spain, Sieverd. & N.C. Schenck, Mycologia 76 (4): 686 (1984)

MycoBank MB 105884

Synonym: *Ambispora appendicula* (Spain, Sieverd. & N.C. Schenck) C. Walker, Mycol. Res. 112 (3): 298 (2008)

MycoBank MB 511420

Synonym: *Appendicispora appendicula* (Spain, Sieverd. & N.C. Schenck) Spain, Oehl & Sieverd., Mycotaxon 97: 170 (2006)

MycoBank MB 510320

Other species: *Appendiculaspora callosa*, *Appendiculaspora jimgerdemannii*, *Appendiculaspora leptoticha Appendiculaspora callosa* (Sieverd.) Sieverd., G.A. Silva & Oehl, comb. nov., Figs. 18-24 MycoBank MB 862215

Basionym: *Glomus callosum* Sieverd., Angew. Bot. 62 (5-6): 374 (1988)

MycoBank MB 134246

Synonym: *Ambispora callosa* (Sieverd.) C. Walker, Vestberg & A. Schüßler, Mycol. Res. 111 (2): 148 (2007)

MycoBank MB 510213

Synonym: *Appendicispora callosa* (Sieverd.) C. Walker, Vestberg & A. Schüßler, Mycol. Res. 111 (3): 254 (2007)

MycoBank MB 510504

Emended description: Sporocarps unknown; spore formation singly upon a short appendix that arises laterally on the tapering hyphal neck of a sporiferous saccule (acaulo-appendiculosporoid), and glomoid spores formed terminally on hyphae, occurring singly or in loose clusters on extra- or intraradical hyphae (glomoid-appendiculosporoid). Acaulo-appendiculosporoid spores 200-280 μm, hyaline when young, becoming subhyaline to light creamy with age, OW 4-8 μm, OWL1 hyaline, 0.8-1.3 μm, evanescent, crowded with minute (spiny-)warts (about 0.5 high and 0.5-0.7 μm broad) on the outer spore surface; OWL2 hyaline to subhyaline, becoming light creamy to creamy with age, 3.5-7.0 μm, MW hyaline, 4.0-7.0 μm in total, comprising two layers: the outer layer MWL1, 2.0-4.0 μm thick, continuous with the thick, hyaline layer of the appendix and second layer of the connected hypha and sporiferous saccule, and with a convex, alveolate reticulum. These undulations are about 5.0-8.0 μm wide and 1.9-2.6 μm deep. MWL2 is hyaline, 2.0-4.0 μm thick, tightly adherent to MWL1 and thus showing a similar alveolate structure with concave hemispherical depressions on the outer surface that fit into the convex protuberances at the inner surface of MWL1. MWL2 may close the pore of the appendix. MWL1 and MWL2 separate when pressure ruptures the spore. IW 3-6 in total μm; a very thin outer layer IWL1 and a thin inner layer IWL3 appear to adhere to the finely laminated middle layer IWL2 in water mounted specimens, but neither IWL1 nor IWL3 are usually detected in specimens mounted in PVLG. The pedicel (appendix) arises laterally from the hyphal neck of a sporiferous saccule, is often persistent, and resembles a subtending hypha, 30-100 μm long, 20-40 μm wide, cylindric or funnel-shaped tapering to 10-25 μm at the distal end from the spore. Glomoid spores are hyaline to subhyaline, 220-280 μm, with one spore wall (SW), crowded with minute (spiny-)warts (about 0.5 high and 0.5-0.7 μm broad) on the outer spore surface consisting of the mucilaginous SWL1, about 0.5-1.5 μm thick, frequently with adhering debris on the outer surface; SWL2 hyaline, about 2.5-11 μm thick, laminated. Subtending hypha cylindric to slightly funnel shaped, 22-35 diam at the spore base tapering to 10-12 μm within 10-120 μm distance. The pore usually is open, 8-21 μm wide, but sometimes a thin septum deriving from the inner hyphal wall layer is seen, which can be found even at a distance > 100 from the spore base in the transition to the mycelia hyphae.

*Appendiculaspora jimgerdemannii* (Spain, Oehl & Sieverd.) Sieverd., G.A. Silva & Oehl, comb. nov., Figs 16-17

MycoBank MB 862216

Basionym: *Appendicispora jimgerdemannii* Spain, Oehl & Sieverd., Mycotaxon 97: 174 (2006) MycoBank MB 510322

Synonym: *Ambispora jimgerdemannii* (Spain, Oehl & Sieverd.) C. Walker, Mycol. Res. 112 (3): 298 (2008)

MycoBank MB 511417

*Appendiculaspora leptoticha* (N.C. Schenck & G.S. Sm.) Sieverd., G.A. Silva & Oehl, comb. nov.

MycoBank MB 862217

Basionym: *Glomus leptotichum* N.C. Schenck & G.S. Sm., Mycologia 74 (1): 82 (1982)

MycoBank MB 110705

Synonym: *Ambispora leptoticha* (N.C. Schenck & G.S. Sm.) C. Walker, Vestberg & A. Schüßler, Mycol. Res. 111 (2): 148 (2007)

MycoBank MB 510212

Synonym: *Appendicispora leptoticha* (N.C. Schenck & G.S. Sm.) C. Walker, Vestberg & A. Schüßler, Mycol. Res. 111(3): 255 (2007)

MycoBank MB 510501

***Ephemerapareta*** gen. nov. G.A. Silva, Palenz., Sieverd. & Oehl, Figs 25-27 MycoBank MB 862218

Description: Sporocarp formation unknown. Species bimorphic differentiating acaulosporoid and glomoid spores, with spore wall organizations that are typical for Ambisporaceae. Acaulosporoid spores form singly in soils on a short pedicel that branches laterally from the neck of a sporiferous saccule. Glomoid spores are formed singly or in clusters. Sporiferous saccules are hyaline to subhyaline, globose to subglobose, 100-200 µm. The saccule wall is bi-layered, consisting of a subhyaline to light yellow or light creamy, rapidly degrading, evanescent outer layer and a hyaline to subhyaline, semi-persistent inner layer (SWL2; 1.1-2.5 µm thick). The hyphal neck of the saccule is 15-30 µm wide at the spore base and tapers to 5-10 µm within 200-350 µm. The sporiferous saccules often detach from the mature spores while the pedicel often persists on the spore. Pedicels of acaulosporoid spores branch from the saccule neck about 60-160 µm from the saccule terminus. The acaulosporoid spores are 90-135 x 100-150 µm, hyaline to white to white-yellow in water and generally become ochre yellow with age, and when mounted in PVLG. They are white when the outer wall has sloughed from the spore. The spores have three walls: a three-layered outer wall (OW), a bi-layered middle wall (MW) and a three-layered inner wall (IW). The OW is continuous with the undifferentiated outer wall layer of the pedicel and with SWL1 of the saccule wall, and evanescent, short-lived. Middle wall is bi-layered, hyaline, in total 2.4-4.0 µm thick and semi-permanent to regularly evanescent. MWL1 is 1.5-3.0 µm thick and continuous with the inner layer of the pedicel and with SWL2 of the saccule wall. MWL2 is 1.0-2.0 µm thick, tightly adhered to MWL1 and forms the final pore closure of the pedicel at the spore base by continuous wall material. Under slight pressure on the cover slide, both layers easily disaggregate into several to many small (about 5-15 x 5-10 µm), irregular pieces, indicating the fragile nature of the middle wall. Inner wall is hyaline, about 2-5 µm thick and has three wall layers that generally are easily detectable. The IW forms de novo, presumably after the spore pore has been closed by the inner layer of the middle wall (MWL2). Second layer IWL2 is finely laminate and 2.0-3.5 µm thick. IW functions as germinal wall as the germ structure, and subsequently germ tubes emerge from this wall through an initial germ pore (gp). Pedicel of acaulosporoid spores is 10-25 µm long and 10-25 µm wide at the spore base tapering to 7-15 µm from the spore base. Sometimes two wall layers on the pedicel were observed that are continuous with OW and MWL1 of the spore wall and with the saccule wall, but often the OW sloughs. In other cases, the pedicel disappears from the spore base leaving a collar formed by the OW. Sometimes one to two septa arise in the pedicel from MWL1. Germination: the main germ tube generally grows straight through the pedicel and the collar of the OW if they remain present on the germinating spores. A lobed germination structure (gs) emerges in germinating spores consisting of a few (3-6), 5-15 µm long lobes and 1-5 germ tubes arising between the lobes. Glomoid spores are hyaline to subhyaline and have two wall layers that are continuous with the two wall layers of the mycelia hyphae. The outer layer (SWL1) is evanescent, hyaline to subhyaline in water. It may become light yellow to light creamy with age and when mounted in PVLG. The inner layer (SWL2) is persistent, hyaline to subhyaline and finely laminate. The SH is cylindrical to slightly funnel-shaped; spore pore sometimes is closed by a septum, but generally the spore pore appears to be open. Germinating glomoid spores were not found. Formation of vesicular arbuscular mycorrhiza with mycorrhizal structures that stain pale blue in trypan blue.

Etymology: Ephmera-(evanescent, short-lived), -pareta (walls), as two of three spore walls are considered to be short-lived, rapidly degrading, not permanent.

Type species: *Ephemerapareta granatensis* (J. Palenzuela, N. Ferrol & Oehl) Oehl, Palenz., Sieverd. & G.A. Silva, comb. nov.

MycoBank MB 862219

Basionym: *Ambispora granatensis* J. Palenzuela, N. Ferrol & Oehl, Mycologia 103 (2): 334 (2011)

MycoBank MB 513528

***Pelotaspora*** Oehl, G.A. Silva, Palenz. & Sieverd., gen. nov., Figs 28-40 MycoBank MB 862220

Description: Sporocarp formation unknown; spores resemble ambisporoid-acaulosporoid spores, but apparently are formed singly on short hyphae. The spores are globose (112-187 µm in diameter) to oval to ovoid to rarely irregular, 100-250 × 90-200 µm, hyaline or white to yellow or creamy brown to brown. The spores have three walls: a bi-(to triple-)-layered outer wall (OW), a bi-layered middle wall (MW) and a three-layered inner wall (IW). OW is evanescent to semi-persistent. The structural second layer OWL2 is 1.0-6 µm thick and generally stains reddish purple in Melzer’s reagent. The pore collar at the spore base is 7-40 µm wide. MW is bi-layered, hyaline and in total 2.5-5.0 µm thick. It forms a conspicuous reticulum with 5-7 sided, large pits (15-30 x 12-25 µm diam) that are framed by 3-8 µm broad ridges giving the spore a football-like appearance. MWL1 may stain yellow in Melzer’s reagent, at least in the first hours after placing. IW is hyaline, in total 3-5 µm thick. The IW may function as germinal wall as related to many Glomeromycota species with de novo forming IW during spore development.

Etymology: Pelota-(ball) and -spora (spora), referring to the football-like structure of the spores Type pecies: *Pelotaspora reticulata* (Oehl & Sieverd.) Oehl, Palenz., G.A. Silva & Sieverd., comb. nov., Figs 28-31

MycoBank MB 862221

Basionym: *Ambispora reticulata* Oehl & Sieverd., J. Appl. Bot. Food Quality 85 (2): 130 (2012) MycoBank MB 800269

Other species: *Pelotaspora austrolatina*

***Pelotaspora austrolatina***. Oehl, C. Castillo, Palenz., I.C. Sánchez & G.A. Silva, sp. nov., Figs 32-40

MycoBank MB 862222

Type: Holotype 56-5601 (Z+ZT: ZT Myc 15115) isolated from the rhizosphere of *Triticum aestivum* (community Gorbea, Araucanía region, Chile) at about 590 m a.s.l. (39°08’ South, 72°38’ West). Isotype specimens from the same field samples (56-5602 to 56-5610), deposited at Z+ZT (ZT Myc 15116); isotypes (56-5611 to 56-5613) deposited at HURM (HURM 83558-83560); paratypes from the rhizosphere of *Triticum aestivum* (community Curacautín, Araucanía region, Chile) deposited at Z+ZT (ZT Myc 15117). Collection date: 15.9.2011.

Collectors: Claudia Castillo, Victor San Martín, Fritz Oehl.

Etymology austro-(southern), and -latina (latino) referring to the first findings in the South of Brazil and Chile.

Description: Sporocarp formation unknown; spores resemble ambisporoid-acaulosporoid spores but are formed singly on short hyphae. The spores are globose (112-187 µm in diameter) to oval to ovoid to rarely irregular, 125-195(−245) × (95-)102-155(−180) µm, hyaline to white to yellow-white, sometimes becoming light ochre-yellow with age or with time when mounted in PVLG. They have three walls: a bi-(to triple-)-layered outer wall (OW), a bi-layered middle wall (MW) and a three-layered inner wall (IW).

*Outer wall* generally consists of two to three layers. The outer layer (OWL1) is hyaline to white to white yellow, 0.8-2.1 µm thick and evanescent to semi-persistent. The structural second layer OWL2 is 1.0-2.5 µm thick. It generally stains reddish purple in Melzer’s reagent. The third layer OWL3 is difficult to detect as it is thin (< 0.6 µm) and tightly adherent to OWL2, and is generally hidden by the outer layer of the middle wall (MWL1).

*Middle wall* is bi-layered, hyaline and in total 2.8-4.4 µm thick. It forms a conspicuous reticulum with 5-6 sided, large pits (15-27 x 12-24 µm diam) that are framed by (2.5-)4.5-7.9 µm broad ridges giving the spore a football-like appearance. MWL1 is 1.4-2.5 µm thick. MWL2 is 1.0-2.5 µm thick, tightly adherent to MWL1 and regularly stains yellow in Melzer’s reagent, at least in the first hours after placing.

*Inner wall* is hyaline, in total 3.0-4.5 µm thick and has three wall layers. The outer layer IWL1 is 0.5-1.2 µm, tightly adherent to the central layer IWL2. Second layer IWL2 is finely laminate and 2.0-2.4 µm thick. Innermost layer IWL3 is 0.5-0.9 µm thick and also tightly adherent to IWL2. IWL3 sometimes separates slightly from IWL2 and often shows several thin folds especially when pressure is applied to the cover slip.

*Pedicel*-like subtending hyphae of these pelotasporoid spores at the spore base is 7.5-25(−38) µm wide. Formation of mycorrhizal structures is so far unknown.

*Distribution.* So far, *Pe. austrolatina* was isolated from two field sites in the Araucanía region of Southern Chile subjected to winter wheat production, and from a viticultural field site in Southern Brazil. It was also found in a grassland and a deciduous forest in the ‘de Los Rios’ region close to Valdivia, Southern Chile. It co-occurred with other AM fungi like *Acaulospora punctata*, *Entrophospora claroidea*, *En.infrequens*, *Funneliformis mosseae* and *Septoglomus constrictum* at the San Pablo de Tregua Experimental Station of Valdivia (Castillo et al., 2006, Oehl et al., 2011c), with AM fungi like *Ac. sieverdingii* (Oehl et al., 2011d), *Paraglomus occultum* and *Rhizoglomus invermaium* in Gorbea and Curacautin, and, among others, with several *Acaulospora* species, *S. constrictum* and *Simiglomus hoi*, *Am. gerdemannii*, *Scutellospora calospora* and *Fuscutata heterogama* in Caxias (RS, Brazil).

### 3.4. Morphological identification key to the Ambisporaceae species

Here, the first morphological identification key is presented for the species attributed to the fungal family Ambisporaceae.

1 Spores with three walls, generally formed on sporiferous saccules, generally on a lateral branch of the saccule neck (ambisporoid-acaulosporoid), rarely on subtending hyphae (triple-walled-glomoid, resembling ambisporoid-acaulosporoid morph) 2

1 Mono-walled spores formed on subtending hyphae (ambisporoid-glomoid morph) 11

2 Spores not formed on sporiferous saccules (triple-walled-glomoid: football like, reticulate ornamentation on the outer surface of MW: *Pelotaspora* 3

2 Spores formed on sporiferous saccules, generally on a lateral branch of the saccule neck (ambisporoid-acaulosporoid sensu lato) 4

3 Spores hyaline to white to rarely pale yellow, 125-195(−245) × (95-)102-155(−180) µm, MW with a diagnostic reticulate ornamentation on the surface consisting of 5-6 sided pits, 15-27 x 12-24 µm framed by (2.5-)4.5-7.9 µm broad ridges. *Pelotaspora austrolatina*

3 Spores yellow-brown to brown, 87-131 x 125-150 μm MW with reticulate outer surface with irregular triagonal to octagonal (usually tetra- to hexagonal) pits that are surrounded by ridges. *Pelotaspora reticulata*

4 Spores with two hyaline, smooth inner spore walls (MW & IW) below the white to creamy or brown outer wall (OW) 5

4 Spores with alveolate MW and smooth IW below the white to creamy, brown or orange-brown OW (*Appendiculaspora*) 6

5 OW and MW (semi-)permanent to evanescent, MW not persistent, often dissolving in liquid mountants, upon pressure on the cover-slide; spores 90-150 μm diam and hyaline to white to pale yellow. Outer surface of OW rough and crowded with papillae, which might become difficult to detect within a few hours in lactic acid-based mountings, but they are clearly visible in water. The structural central wall layer of the outer wall is only 0.8-1.5 μm thick *Ephemerapareta granatensis* (ambisporoid-acaulosporoid morph)

5 MW permanent, cracking in regular to irregular plate-like pieces under pressure on the cover-slide (*Ambispora*) 9

6 OW without surface ornamentation 7

6 OW with surface ornamentation 8

7 Middle wall with alveolate ornamentation. Spores (170-)250(−390) μm diam, white-opaque when young, becoming dull yellow-cream to yellow to yellowish-brown or brown, rarely orange-tan; main layer OWL2 firm and difficult to break on young spores, turns orange-red in Melzer’s reagent, but with age becoming less rigid and staining less red with Melzer’s reagent. *Appendiculospora appendicula* (acaulo-appendiculosporoid morph)

7 Spores 160-240 µm, cream when young, becoming orange-brown to dark orange-brown, OWL2 staining dark orange-brown to red-brown in Melzer’s reagent. *Appendiculaspora leptoticha* (acaulo-appendiculosporoid morph)

8 OW with minute (spiny-)warts on the outer spore surface (OWL1), 200-280 μm, hyaline when young, becoming subhyaline to creamy with age, OW 4-8 μm, MW 4-7 μm, IW 3-6 μm. *Ap. callosa* (acaulo-appendiculosporoid morph)

8 OW with labyrinthiforme ornamentation on the outer surface of OWL2; Spores 200-250 μm, brown at maturity; the main layer OWL2 is brown in color, becoming brittle with age. It is approximately (8-)10-14 μm thick showing cerebriform folds over the laminated wall layer that is 1-1.5 μm thick. The cerebriform ridges are up to (6-)10-12 μm high and 4-6 μm broad and 1-3 μm apart from each other; the outer spore wall does not react with Melzer’s reagent. *Appendiculaspora jimgerdemannii* (acaulo-appendiculosporoid morph)

9 Spores < 100 μm are formed on a pedicel branching from the neck of a sporiferous saccule. They are hyaline to pale yellow in color, globose, 62-95 μm in diameter, to subglobose to oval, 59-88 × 69-100(−118) μm, and with three spore walls. Crowded irregular pustules, 2.4-7.0(−10.0) μm high and 4.2-9.8(−15.0) μm wide, are formed on the surface of the outer wall. *Ambispora brasiliensis* (acaulo-appendiculosporoid morph)

9 Spores > 100 μm 10

10 OW not staining purple in Melzer’s reagent 11

10 OW staining rust-brown to red or purple in Melzer’s reagent; spores hyaline to pale ochraceous, to pyriform, 124-201 x 134-201 μm, *Ambispora fennica* (ambisporoid-acaulosporoid morph)

11 MW regularly cracking into rectangular pieces under pressure on the cover slide; spores 160-310 μm, hyaline to pale yellow-brown *Ambispora gerdemannii* (ambisporoid-acaulosporoid morph)

11 Spores MW cracking into irregular pieces under pressure on the cover slide, 100-220 μm, hyaline to pale yellow-brown *Ambispora nicolsonii* (ambisporoid-acaulosporoid morph)

12 Spores without ornamentation on outer surface 13

12 Spores with surface ornamentation 17

13 Mature spores hyaline to subhyaline, to pale yellow 14

13 Mature spores yellow-brown to dark brown, 100-210 x 80-150 μm *Ambispora fecundispora* (ambisporoid-glomoid morph)

14 Spores regularly < 70 μm 15

14 Spores regularly > 70 μm 16

15 Spores 25-30 μm, hyaline to subhyaline, bi-layered *Ambispora brasiliensis* (ambisporoid-glomoid morph)

15 Spores (25)40-70 μm, hyaline to pale yellow, bi-layered *Ephemerapareta granatensis* (ambisporoid-glomoid morph)

16 Spores 40-130 μm diam, hyaline, with a mucilaginous outer layer SWL1, about 0.8-1.0 μm thick, frequently with adhering debris on the outer surface, and a permanent and laminate SWL2, 1.5-4.0 μm thick. *Ambispora fennica* or *Ambispora gerdemannii* (ambisporoid-glomoid morph)

16 Spores 120-240(−280) μm, hyaline to subhyaline, with a mucilaginous SWL1, 1.5-2.5 μm thick, frequently with adhering debris on the outer surface, and a permanent and laminate SWL2, 2-8(−12) μm thick *Appendiculaspora appendicula* (ambisporoid-glomoid morph)

17 Spores subhyaline, 220-280 μm, spore surface crowded with minute warts *Appendiculaspora callosa* (ambisporoid-glomoid morph)

17 Spores hyaline, when young; mature spores light yellow; 150-265 x 100-235 μm; spore surface with an alveolate reticulum of shallow ridges, 0.5-1 wide *Appendiculospora leptoticha* (ambisporoid-glomoid morph)

## Discussion

Our new phylogenetic analyses indicate three different clades in Ambisporaceae (A, B, C; *Ambispora*, *Appendiculaspora*, and *Ephemerapareta*, respectively). It was already demonstrated for Glomerales that clades that differ by ≥ 10% MI, according to the partial nrDNA gene (Corazon-Guivin et al., 2019; Silva et al., 2023; Silva et al., 2025) can be considered different genera. *Ambispora* has 89% MI with *Appendiculaspora*, and the tree topology place, with strong support, *Ambispora* (sensu stricto) in a single clade well separated from the other two clades. *Ephemerapareta granatensis* has been sequenced just to their SSU and ITS nrDNA, however the Blastn analysis of the ITS sequences from this species did not show a relevant MI with any other clade of the family. *Ambispora* clusters together with *Appendiculaspora*, but there is no support to place these genera in the same group in the SSU tree (Fig. 3). The SSU-ITS-LSU and ITS trees (Figs. 1 and 2) have no support for ML-FBP, and a low support for ML-TBE in the ITS tree (82%) to place these genera together. In the ITS and SSU trees performed by Walker et al. (2007), Ambisporaceae is clearly separated in two clades (the first one with *Am. fennica* and the second one with *Am. gerdemannii*, *Ap. leptoticha* and *Ap. callosa*. At that time *Ep. granatensis* (Palenzuela et al., 2011) was not published and there were no sequences from *Ap. appendicula* so far available.

In the tree, generated by Esmaeilzadeh-Salestani et al. (2025), also a separation of *Ambispora* (*sensu lato*) was observed into two clades (the first one with *Ap. appendicula* and *Ap. leptoticha* and the second one with *Am. fennica* and *Am. gerdemannii*). In our ITS tree (Fig. 2), distinct isolates of *Am. gerdemannii* are placed in different clades (together with *Ap. appendicula/leptoticha*/*callosa* clade or *Am. fennica* clade), corroborating the results of both papers, and indicating a mistake with some sequences attributed to *Am. gerdemannii* in the NCBI. In the SSU tree (Fig. 3), sequences from the isolate MAFF520057 are attributed to *Ap. callosa* and also to *Ap. leptoticha*. In this case we believe that the culture presents both species, considering that there are sequences of this isolate in two different clades. The isolate MAFF520058 is attributed to *Ap. leptoticha*, but according to our phylogenetic analyses, possibly this isolate represents in fact *Ap. callosa*. One isolate (n8_9) attributed to *A. gerdemannii* was sequenced for the entire SSU+ITS+LSU nrDNA fragment, while the isolate AU215 was sequenced just for the ITS region, and LSU nrDNA. In Fig. 2, it is possible to see sequences from two isolates of *Am. gerdemannii* in different clades (‘A’ and ‘B’). The reference INVAM culture for *Am. gerdemannii* (MT106) was sequenced just for the LSU nrDNA and is placed in the clade ‘A’, while the *Am. gerdemannii* - isolate AU215 group in the clade B (data not shown), indicating a probable mistake with sequences attributed to *Am. gerdemannii* AU215.

The clear results obtained for the three major clades of the Ambisporaceae in the phylogenetic analyses requested for advanced morphological separations. Morphologically, four major groups had been recognized in the Ambisporaceae: i) bi-morphic species with an ambisporoid-acaulosporoid and an ambisporoid-glomoid morph, ii) species, for which so far only an ambisporoid-acaulosporoid morph has been detected, iii) species for which hitherto only an ambisporoid-glomoid morph has been known, and iv) species with triple walled glomoid-like spores formed on hyphae, whose spores resemble the ambisporoid-acaulosporoid, but their formation on sporiferous saccules has not yet been observed so far. All these species were summarized before within one genus, which is *Ambispora*.

These major morphological similarities respective differences are based on the spore wall composition of the ambisporoid-acaulosporoid morph, mainly based on the structure of the central wall, as presented above: i) *Am. fennica*, *Am. gerdemannii*, *Am. nicolsonii* and *Am. brasiliensis* have a smooth, permanent central spore wall and representing the revised genus *Ambispora*, ii) the central wall of *Ap. appendicula*, *Ap. callosa*, *Ap. leptoticha and Ap. jimgerdemannii* is also permanent, but conspicuously alveolate (new genus *Appendiculaspora*), and iii) the central wall of *Ep. granatensis* is smooth, but easily degraded, thus rather short-lived and not permanent, but evanescent (new genus *Ephemerapareta*), when compared to species of the clades A and B. On the other hand, the ambisporoid-glomoid morph so far cannot be used for genus identification, since besides spore size, hardly a character can be found, which might separate clearly even on the species level.

For two additional species, *Pe*. *reticulata* and *Pe. austrolatina*, the molecular analyses failed so far. Their spores have also triple-walled spores, and they resemble those triple-walled ambisporoid-acaulosporoid spores of *Ambispora*, *Appendiculaspora* and *Ephermerapareta*, but apparently these spores may not be formed on sporiferous saccules. These species have a diagnostic reticulate middle wall with 4-8 sided, but generally 5-6 sided, large pits that are formed by broad ridges giving the spore a football-like morphological appearance. These species should no longer be counted within the newly organized *Ambispora*, or one of the two new genera (*Appendiculaspora* & *Ephemerapareta*) of the Ambisporaceae, due to their clear diagnostic features. Thus, we here decided to attribute them to the new genus *Pelotaspora*.

Future studies are needed to show their phylogeny, exact spore formation, their intraradical and extraradical fungal structures and germination precisely.

## Acknowledgments

Daniele Magna Azevedo de Assis and Thays Gabrielle Lins de Oliveira thanks the Fundação de Amparo à Ciência e Tecnologia do Estado de Pernambuco (FACEPE) for providing fellowship. Gladstone A. Silva has a fellowship from the Conselho Nacional de Desenvolvimento Científico e Tecnológico (CNPq) (Proc. 312606/2022-2, approval date: 16 February 2023).

## Conflict of interest

No potential conflict of interest was reported by the authors.

**Spreadsheet S1**. GenBank accession numbers for the sequences used in this study.

## Notes

### Competing Interest Statement

The authors have declared no competing interest.

