## Supplementary material for "Revision of Ambisporaceae, with three new genera and one new species and a morphological identification key for all the species currently attributed to this family": Spreadsheet S1

**Spreadsheet S1**. GenBank accession numbers for the sequences used in this study.

| Species name in NCBI | ID | nrDNA region | Country | GenBank accession numbers (nrDNA) |
|---|---|---|---|---|
| *Ambispora fennica* | Att200-23 - ex-type | SSU+ITS+LSU | Finland | FN547535 |
| *Ambispora fennica* | Att200-23 - ex-type | SSU+ITS+LSU | Finland | FN547536 |
| *Ambispora fennica* | Att200-23 - ex-type | SSU+ITS+LSU | Finland | FN547537 |
| *Ambispora fennica* | Att200-23 - ex-type | SSU+ITS+LSU | Finland | FN547538 |
| *Ambispora fennica* | Att200-23 - ex-type | SSU+ITS+LSU | Finland | FN547539 |
| *Ambispora fennica* | Att200-23 - ex-type | SSU+ITS+LSU | Finland | FN547540 |
| *Ambispora fennica* | Att200-23 - ex-type | SSU+ITS+LSU | Finland | FN547541 |
| *Ambispora fennica* | Att200-23 - ex-type | SSU+ITS+LSU | Finland | FN547542 |
| *Ambispora fennica* | Att200-23 - ex-type | SSU+ITS+LSU | Finland | FN547543 |
| *Ambispora fennica* | Att200-23 - ex-type | SSU+ITS+LSU | Finland | FN547544 |
| *Ambispora fennica* | Att200-23 - ex-type | SSU+ITS+LSU | Finland | FN547545 |
| *Ambispora fennica* | Att200-23 - ex-type | SSU+ITS+LSU | Finland | FN547546 |
| *Ambispora fennica* | Att200-23 - ex-type | SSU+ITS+LSU | Finland | FR750157 |
| *Ambispora fennica* | Att200-21 | SSU | Finland | AM268192 |
| *Ambispora fennica* | Att200-21 | SSU | Finland | AM268193 |
| *Ambispora fennica* | Att200-21 | SSU | Finland | AM268194 |
| *Ambispora fennica* | Att200-23 - ex-type | SSU | Finland | AM268196 |
| *Ambispora gerdemannii* | n8_9 | SSU+ITS+LSU | Not available | JF439210 |
| *Ambispora gerdemannii* | AU215 | ITS | Australia | AJ012111 |
| *Ambispora gerdemannii* | AU215 | ITS | Australia | AM743187 |
| *Appendiculaspora appendicula* | Att1235-1 | SSU+ITS+LSU | Brazil | FN547524 |
| *Appendiculaspora appendicula* | Att1235-1 | SSU+ITS+LSU | Brazil | FN547525 |
| *Appendiculaspora appendicula* | Att1235-1 | SSU+ITS+LSU | Brazil | FN547527 |
| *Appendiculaspora appendicula* | Att1235-1 | SSU+ITS+LSU | Brazil | FN547528 |
| *Appendiculaspora appendicula* | Att1235-1 | SSU+ITS+LSU | Brazil | FN547529 |
| *Appendiculaspora appendicula* | Att1235-1 | SSU+ITS+LSU | Brazil | FN547530 |
| *Appendiculaspora appendicula* | Att1235-1 | SSU+ITS+LSU | Brazil | FN547531 |
| *Appendiculaspora appendicula* | Att1235-1 | SSU+ITS+LSU | Brazil | FN547532 |
| *Appendiculaspora appendicula* | Att1235-1 | SSU+ITS+LSU | Brazil | FN547533 |
| *Appendiculaspora appendicula* | Att1235-1 | SSU+ITS+LSU | Brazil | FN547534 |
| *Appendiculaspora appendicula* | MACG1 | SSU+ITS+LSU | Peru | OM214461 |

| | | | | |
|---|---|---|---|---|
| *Appendiculaspora appendicula* | SA19006 | SSU | Not available | MN784846 |
| *Appendiculaspora callosa* | JA116 | SSU+ITS+LSU | Japan | PX215118 |
| *Appendiculaspora callosa* | MAFF520084 | SSU+ITS+LSU | Japan | PX215120 |
| *Appendiculaspora callosa* | MAFF520084 | SSU+ITS+LSU | Japan | PX215121 |
| *Appendiculaspora callosa* | JA116 | SSU+ITS+LSU | Japan | PX215122 |
| *Appendiculaspora callosa* | MAFF520084 | SSU+ITS+LSU | Japan | PX215125 |
| *Appendiculaspora callosa* | JA116 | SSU+ITS+LSU | Japan | PX215126 |
| *Appendiculaspora callosa* | MAFF520084 | SSU+ITS+LSU | Japan | PX215128 |
| *Appendiculaspora callosa* | MAFF520084 | SSU+ITS+LSU | Japan | PX215129 |
| *Appendiculaspora callosa* | MAFF520084 | SSU+ITS+LSU | Japan | PX215131 |
| *Appendiculaspora callosa* | JA116 | SSU+ITS+LSU | Japan | PX215132 |
| *Appendiculaspora callosa* | MAFF520057 | SSU+ITS+LSU | Japan | PX215133 |
| *Appendiculaspora callosa* | MAFF520057 | SSU+ITS+LSU | Japan | PX215134 |
| *Appendiculaspora callosa* | MAFF520057 | SSU+ITS+LSU | Japan | PX215135 |
| *Appendiculaspora callosa* | JA116 | SSU+ITS+LSU | Japan | PX215136 |
| *Appendiculaspora callosa* | JA116 | SSU+ITS+LSU | Japan | PX215137 |
| *Appendiculaspora callosa* | Att1321-4 | ITS | Japan | AB259842 |
| *Appendiculaspora callosa* | MAFF520073 | ITS | Japan | AB259846 |
| *Appendiculaspora leptoticha* | MAFF520090 | SSU+ITS+LSU | Japan | PX215138 |
| *Appendiculaspora leptoticha* | ON205A | SSU+ITS+LSU | Canada | PX215139 |
| *Appendiculaspora leptoticha* | ON205A | SSU+ITS+LSU | Canada | PX215140 |
| *Appendiculaspora leptoticha* | ON205A | SSU+ITS+LSU | Canada | PX215141 |
| *Appendiculaspora leptoticha* | MAFF520055 | ITS | Japan | AB048630 |
| *Appendiculaspora leptoticha* | NC176 | ITS | USA | AJ012109 |
| *Appendiculaspora leptoticha* | FL130 | ITS | USA | AJ012201 |
| *Appendiculaspora leptoticha* | MAFF520055 | SSU | Japan | AB047302 |
| *Appendiculaspora leptoticha* | MAFF520055 | SSU | Japan | AB047303 |
| *Appendiculaspora leptoticha* | MAFF520055 | SSU | Japan | AB047304 |
| *Appendiculaspora leptoticha* | MAFF520057 | SSU | Japan | AB047306 |
| *Appendiculaspora leptoticha* | MAFF520057 | SSU | Japan | AB047307 |
| *Appendiculaspora leptoticha* | MAFF520058 | SSU | Japan | AB047308 |
| *Appendiculaspora leptoticha* | MAFF520058 | SSU | Japan | AB047309 |
| *Appendiculaspora leptoticha* | NC176 | SSU | USA | AJ006466 |

| | | | | |
|---|---|---|---|---|
| *Appendiculaspora leptoticha* | NC176 | SSU | USA | AJ301861 |
| *Archaeospora trappei* | Att178-3 | SSU+ITS+LSU | UK | FR750034 |
| *Archaeospora trappei* | Att178-3 | SSU+ITS+LSU | UK | FR750035 |
| *Archaeospora trappei* | Att178-3 | SSU+ITS+LSU | UK | FR750036 |
| *Archaeospora trappei* | Att178-3 | SSU+ITS+LSU | UK | FR750037 |
| *Archaeospora trappei* | Att178-3 | SSU+ITS+LSU | UK | FR750038 |
| *Archaeospora trappei* | NB112 | SSU | Namibia | AJ006800 |
| *Archaeospora trappei* | Att186-1 | SSU | Austria | AM114274 |
| *Archaeospora trappei* | Att186-1 | SSU | Austria | Y17634 |
| *Ephemerapareta granatensis* | ex-type | ITS | Spain | FN820276 |
| *Ephemerapareta granatensis* | ex-type | ITS | Spain | FN820277 |
| *Ephemerapareta granatensis* | ex-type | ITS | Spain | FN820278 |
| *Ephemerapareta granatensis* | ex-type | ITS | Spain | FN820280 |
| *Ephemerapareta granatensis* | ex-type | ITS | Spain | FN820281 |
| *Ephemerapareta granatensis* | ex-type | ITS | Spain | FN820282 |
| *Ephemerapareta granatensis* | ex-type | SSU | Spain | FN820272 |
| *Ephemerapareta granatensis* | ex-type | SSU | Spain | FN820274 |
| *Geosiphon pyriformis* | GEO1 | SSU+LSU | Germany | FM876840 |
| *Geosiphon pyriformis* | GEO1 | SSU+LSU | Germany | FM876841 |
| *Geosiphon pyriformis* | GEO1 | SSU+LSU | Germany | FM876842 |
| *Geosiphon pyriformis* | GEO1 | SSU+LSU | Germany | FM876843 |
| *Geosiphon pyriformis* | GEO1 | SSU+LSU | Germany | FM876844 |
| *Geosiphon pyriformis* | GEO1 | SSU | Germany | AM183923 |
| *Geosiphon pyriformis* | GEO1 | SSU | Germany | AJ276074 |
| *Geosiphon pyriformis* | GEO1 | SSU | Germany | Y15904 |
| *Geosiphon pyriformis* | GEO1 | SSU | Germany | Y15905 |
| *Paraglomus brasilianum* | Att260-8 | SSU+ITS+LSU | Brazil | FR750046 |
| *Paraglomus brasilianum* | Att260-8 | SSU+ITS+LSU | Brazil | FR750047 |
| *Paraglomus occultum* | Att677-4 | SSU | USA | AJ276081 |
| *Paraglomus occultum* | Att677-3 | SSU | USA | AJ276082 |
| *Polonospora polonica* | ex-type | SSU+ITS+LSU | Poland | MZ359654 |
| *Polonospora polonica* | ex-type | SSU+ITS+LSU | Poland | MZ359655 |
| *Polonospora polonica* | ex-type | SSU+ITS+LSU | Poland | MZ359656 |

| | | | | |
|---|---|---|---|---|
| *Polonospora polonica* | ex-type | SSU+ITS+LSU | Poland | MZ359657 |
| *Polonospora polonica* | ex-type | SSU+ITS+LSU | Poland | MZ359658 |
| *Polonospora polonica* | ex-type | SSU+ITS+LSU | Poland | MZ359659 |
